# The co-receptors *Orco* and *Ir8a* are required for coordinated expression of chemosensory genes in the antennae of the yellow fever mosquito, *Aedes aegypti*

**DOI:** 10.1101/2025.04.23.650034

**Authors:** Matthew M. Cooke, Michael S. Chembars, R. Jason Pitts

## Abstract

Olfaction has been extensively studied in the yellow fever mosquito, *Aedes aegypti*. This species uses its sense of smell to find blood hosts and other resources, contributing to its impact as a vector for human pathogens. Two major families of protein-coding genes, the odorant receptors (*Ors*) and the ionotropic receptors (*Irs*), provide the mosquito with sensitivities to distinct classes of volatile compounds in the antennae. Individual tuning receptors in both families require co-receptors for functionality, *Orco* for all *Ors*, and *Ir8a* for many *Irs*, especially ones that are involved in carboxylic acid detection. In *Drosophila melanogaster*, disruptions of *Orco* or *Ir8a* impair receptor function, tuning receptor expression, and membrane localization, leading to general anosmia. We reasoned that *Orco* and *Ir8a* might also be important for coordinated chemosensory receptor expression in the antennal sensory neurons of *Ae. aegypti*. To test this, we performed RNAseq and differential expression analysis in wild type versus *Orco*^−/−^ and *Ir8a*^−/−^ mutant adult female antennae. Our analyses revealed *Or* and *Ir* tuning receptors are broadly under-expressed in *Orco*^−/−^ mutants, while a subset of tuning *Irs* are under-expressed in *Ir8a* mutants. Other chemosensory and non-chemosensory genes are also dysregulated in these mutants. Further, we identify differentially expressed transcription factors including homologs of the *Drosophila melanogaster Mip120* gene. These data suggest a previously unknown pleiotropic role for the *Orco* and *Ir8a* co-receptors in the coordination of expression of chemosensory receptors within the antennae of *Ae. aegypti* by participating in a feedback loop involving *amos* and members of the *MMB/dREAM* complex.

**Simple Summary:** Mosquitoes have an exquisite olfactory system, with which they locate bloodmeal hosts, nectar sources, and sites for egg-laying. Elucidating the mechanisms that underlie the perception of and response to chemicals in the environment is crucial for our understanding of the biology of mosquitoes that are vectors of many deadly pathogens. Olfaction is primarily mediated by large families of odorant receptors and ionotropic receptors, collectively encoding more than 200 tuning receptors, each of which recognizes one or more volatile odorants. As ligand-gated ion channels, tuning receptors of each family form complexes with the Odorant receptor co-receptor (*Orco*), or one of few Ionotropic receptor co-receptors (*Irco*), respectively. In this study, we evaluated the hypothesis that co-receptors are necessary for tuning receptor expression. To this end, we compared antennal transcriptomes of wild type adult female *Aedes aegypti* with *Orco*^−/−^ and *Ir8a*^−/−^ mutant strains. We show that the tuning receptor transcripts in the antennae are broadly dysregulated in both mutants. We discuss two possible explanations for these observations and suggest ways this knowledge can be applied to vector control strategies.

Graphical Abstract

## 1. Introduction

Our understanding of fly behavior, physiology, and olfaction has been largely influenced by studies of the genetic model, *Drosophila melanogaster*, with more recent contributions in the disease vector, *Aedes aegypti* [1–7]. *Ae. aegypti* is primarily responsible for the spread of arboviruses such as Dengue, Yellow fever, Chikungunya, and Zika [8]. Dengue fever alone accounted for an estimated 6.5 million cases, with more than 7300 deaths globally in 2023 [9,10]. These mosquitoes spread pathogens via hematophagous behavior, and they locate their blood hosts predominantly via olfaction [11–13]. Therefore, understanding mosquito olfaction not only answers questions about the basic biology of insects but also informs vector surveillance and control strategies that target host-seeking behavior.

Mosquitoes detect semiochemicals in their environment via an elegant olfactory system [13]. Though the antennae are the major olfactory organs, the maxillary palp, labellum, and tarsi also contribute to olfaction [14]. On the surface of each olfactory organ are hairlike sensilla, which each contain one or more dendrites of olfactory sensory neurons (OSNs) [2]. These neurons characteristically express odorant receptors (*Ors*) or ionotropic receptors (*Irs*), the two major chemosensory receptor families involved in olfaction. Recent analysis has demonstrated co-expression of these classes of chemoreceptors [2]. Gustatory receptors (*Grs*) are a different class of chemosensory receptor that are involved in taste and carbon dioxide detection [15–17]. These three chemoreceptor classes are used in coordination by the insect to detect semiochemicals and thus locate resources in their niche environments, including potential blood hosts [18,19].

*Ors* were the first insect chemoreceptor genes to be discovered and have been extensively studied [2,5,20,21]. *Irs* were discovered more recently but appear to be the more ancient gene family as they are found in the genomes of protostomes [20–22]. Not only are these two classes of receptors understood to be distinct in their evolutionary lineages, but they are also expressed in different types of sensilla and respond to different classes of volatile compounds [23]. Ors typically respond to floral odors including indoles and alcohols, among others [24], while Irs respond predominantly to aldehydes, carboxylic acids, amines, and ketones [25–29]. For this reason, Ors have been implicated in mosquitoes as mediating behaviors such as nectar feeding, while Irs are considered responsible for mediating host-seeking and oviposition behaviors [24,25,30].

Ors and Irs each comprise ligand-gated ion channels, where the cognate volatile compound binds to the receptor and opens the channel [31,32]. For each chemoreceptor family, one or more co-receptors cooperates with a tuning receptor to form a functional ion channel [29].

Stoichiometrically, three odorant receptor co-receptor (Orco) polypeptide subunits, form a complex with a single tuning receptor (Orx) [31,32]. The tuning receptors contribute to the specificity of the ligand-gated ion channel by binding to cognate ligands [24,33]. In contrast, co-receptors for the Irs include Ir8a, Ir25a, Ir76b, and Ir93a [34,35]. These co-receptors function in a similar way to *Orco*, where the Ir co-receptor must be present in complex with a tuning receptor to form a functional channel, although their exact stoichiometry remains unresolved [23]. Ir8a seems to be required for the formation of carboxylic acid-sensitive ion channels that are activated by acetic, lactic, and nonanoic acids [6,27,29].

In *D. melanogaster* and *Ae. aegypti* several studies have demonstrated behavioral deficits in flies lacking the *Orco* or *Ir8a* co-receptors [6,27,36–38]. In *Orco* mutants, Or tuning receptor trafficking from the endoplasmic reticulum to the dendrite surface was impaired [36,39]. Thus, *Orco* may coordinate the subcellular localization of tuning Ors. Similarly, in an *Ir8a* mutant, expression of the *Ir64a* tuning receptor was absent in the sensilla of the antennal sacculus. Instead, low levels of the protein were found in the ER [38]. Notably, not only was transport of *Ir64a* affected in the *Ir8a* mutant, but the overall abundance of *Ir64a* in the antennae was significantly reduced [38]. Another study indicated that, some but not all tuning receptor localization to the cilia was disrupted in an *Ir8a* mutant [40]. Odorant receptor neuron degeneration has also been observed in the maxillary palps of *Orco* mutants [41–43]. Furthermore, a recent study has demonstrated that knockouts of *Orco* and *Ir8a* in *D. melanogaster* lead to disrupted transcription of chemoreceptors, including *Ors* and *Irs* [44].

Genetic disruptions of *Orco* and *Ir8a* in *Ae. aegypti* has produced mutant strains displaying general anosmia [27,37]. Given our understanding of chemoreceptor gene regulation in *D. melanogaster*, we hypothesized that *Orco^−/−^* or *Ir8a^−/−^* disruptions in *Ae. aegypti* might also lead to reduced expression of tuning receptors in the antennae compared to wildtype. The under-representation of these transcripts in mutant antennae could result from dysregulated transcription, a neurodegenerative process, or a combination of the two. To test this prediction, we analyzed the antennal transcriptomes of *Ae. aegypti Orco*^-16^ and *Ir8a*^dsRED/dsRED^ mutant strains [27,37] and compared them to the background wildtype Orlando strain.

## 2. Materials and Methods

### 2.1. Knockout strains

The *Orco*^−/−^ mutant was produced via zinc finger mutagenesis [37]. The mutant has a 16 base pair deletion in the first exon, a frameshift mutation leading to premature stop codons, and a phenotype of general anosmia [37]. The *Ir8a*^−/−^ and the *Ir8a*^attP/attP^ strains were produced via CRISPR/Cas9 mutagenesis and display a phenotype of reduced ability to detect carboxylic acids and reduced host-seeking behavior [27]. The *Ir8a*^−/−^ strain contains a knock-in cassette in the second exon, with a polyubiquitin promoter upstream of the *dsRED* fluorescent gene, leading to the production of *dsRED* in all tissues of the body [27]. The *Ir8a*^attP/attP^ strain contains a 50 bp knock-in at the third exon with an *attP PhiC31* recombination site [27].

### 2.2 Mosquitoes

Mosquitoes were reared under controlled conditions (12:12 LD, 27^℃^, 70% RH, 10% sucrose *ad libitum*), with all strains kept in the same incubator. Eggs were hatched in distilled water (diH_2_O) under vacuum for approximately 1-3 hours. Larvae were reared in clean diH_2_O and were fed a mixture of ground koi fish-food mixed with baker’s yeast *ad libitum*. Larval pan water was filtered regularly to limit the growth of fungus or bacteria. Pupae were separated from larvae and placed into a cup of diH_2_O inside a mesh cage (BugDorm-1; MegaView Sci. Co. Ltd., Taiwan).

Adults were allowed to eclose for two days before pupae cups were moved to a new BugDorm, ensuring all adults had eclosed within a two-day window. Each enclosure contained between 200 and 400 adults of both sexes. Adults were allowed to mature and feed *ad libitum* on 10% sucrose until all mosquitoes were five to seven days post eclosion (DPE). Male and female adults were not separated after eclosion; therefore, we assume that females were mated.

### 2.3 Antennal Dissections

At ZT = 0, the entire cage of mosquitoes was brought to –20°C for approximately one hour, to ensure that all mosquitoes were killed. All surfaces were treated with RNase-away (Thermo Scientific) before dissections. Adult females were then transferred to a chill table at –4°C and antennae were dissected directly into 500µL of TRIzol reagent (Invitrogen) on ice in a 1.5 mL RNase-free microcentrifuge tube. Using surgical forceps, both antennae were resected such that the Johnston’s organ and all thirteen segments of antennae were collected. Each biological replicate consisted of a cage of mosquitoes that was reared to adulthood starting from a separate larval pan. Six replicates were collected for Orlando, four for *Orco*, and five for *Ir8a*. Each sample contained between 100 and 250 pairs of antennae. The tissue was stored at –20°C prior to RNA extraction.

### 2.4 RNA Extractions

RNA was extracted according to the phenol/chloroform extraction method, using TRIzol reagent and substituting 1-bromo-3-chloropropane for chloroform [45]. The tissue was disrupted via repeated cycles of flash freezing and mechanical disruption with an RNase-free pestle. Another 500μl TRIzol reagent was then mixed into each tube. After a 5-minute incubation at room temperature, 200μl of 1-bromo-3-chloropropane was added. The mixture was vortexed vigorously, followed by a 15-minute incubation at room temperature and subsequent centrifugation at 12,000 rcf for 15 minutes at 4°C. The aqueous phase was then carefully pipetted into a new RNase-free tube and an equal volume of 100% isopropanol and 3.0μl of glycogen (5mg/ml Ambion) were added. The samples were stored at –20°C overnight, after which they were centrifuged at 12,000 rcf for 15 minutes at 4°C. The liquid was carefully removed via pipetting without disturbing the pellet of RNA. The pellet was washed with ice-cold 70% ethanol, followed by centrifugation at 7,500 rcf for 10 minutes at 4^℃^. The ethanol was carefully removed via pipetting, without disturbing the pellet, and allowed to dry at room temperature in a biosafety cabinet for ∼5 minutes. The pellet was resuspended in 30 µl of RNase-free H2O, passed through an RNAase-free MicroBiospin-30 column (BioRad), and subjected to SpeedVac (ThermoFisher) evaporation of residual ethanol for 5 minutes. RNA was quantified via NanoDrop (ThermoFisher) and then stored at –80°C. The first samples (Orlando, Orco, and Ir8a samples 1 and 2) were treated with DNaseI during library preparation, leading to low RNA yield. The remaining samples were treated with DNaseI using the Monarch Total RNA Miniprep Kit (NEB #T2010) according to the manufacturer’s instructions. Following DNaseI treatment, samples were again stored at –80°C until they were sent for RNAseq.

### 2.5 RNAseq

RNAseq was outsourced to Psomagen, Inc. Briefly, samples were sent to Psomagen in 30 µl of 10mM Tris. The samples were analyzed via Bioanalyzer to assess sample quality, concentration, and purity. The transcriptome mRNA library was constructed with the Illumina Truseq stranded mRNA library prep kit. RNA was fragmented to 150bp long read length for paired-end sequencing. Sequencing was conducted on the NovaSeq X Plus sequencing system at a 40x coverage. Samples were delivered via an FTP link for download and analysis. Raw reads were submitted the NCBI Sequence Read Archive (SRA) under the accession number PRJNA1249520.

### 2.6 Data analysis

#### 2.6.1 RNAseq data preparation

Raw reads were processed by Trimmomatic to remove the Illumina Universal Adapter and filter out reads with an average quality of <30 across 4 bases using the sliding window trimming approach [46,47]. Trimming and filtering were confirmed via FastQC [48]. A separate annotated transcriptome was constructed for each mutant organism (Supplementary 8,9), with appropriate mutations in the *Orco* or *Ir8a* genes representing alterations in the mutant animals. Salmon was used to perform a pseudoalignment of forward and reverse reads for each sample to the annotated transcriptomes [49]. TxImport was used to transform the output from Salmon into count data and compile it into a matrix for DESeq2 analysis [50]. DESeq2 was performed comparing all three strains to each other, and subsequently, comparisons were made between each mutant strain and Orlando [51,52]. TxImport and DESeq2 were performed in R, and scripts were written with the help of ChatGPT. All R scripts are provided in the Supplementary 10. The workflow for RNAseq processing and analysis is shown in Figure 1.

**Figure 1.**
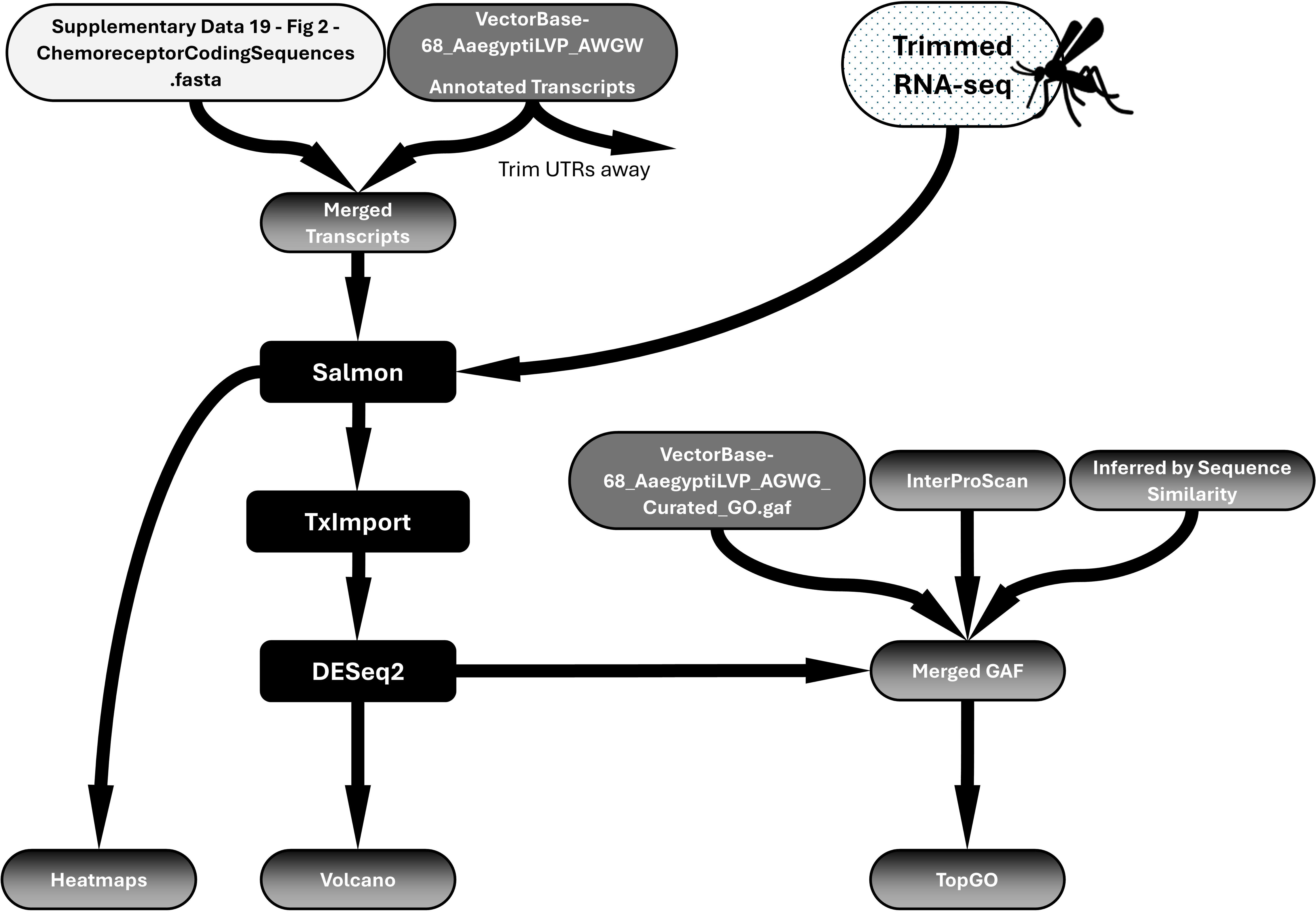
Workflow of RNAseq Differential Expression Analysis. Updated transcriptome files and GAF files were constructed to include current annotations of chemoreceptors. Salmon pseudoalignment was performed to align trimmed reads to the transcriptome. TxImport and DESeq2 analysis were performed to identify differentially expressed genes. Heatmaps, Volcano Plots, and TopGO analysis were performed.

#### 2.6.2 Annotation files preparation

The most comprehensive annotations of chemoreceptors to date were published by [19]. Since these updates were not included in the publicly available L5 *Ae. aegypti* genome annotations, new annotations were constructed to include them. The GFF file describing chemoreceptor genes found in [19] was compared to the VectorBase-68_AaegyptiLVP_AGWG.gff file publicly available on VectorBase using GFFCompare [53]. This tool provided a mapping file, which identified 203 chemoreceptor genes that were common to both GFF files. There were also three chemosensory genes present in the VectorBase GFF, describing two Ors and one Gr, which were not discovered via the GFFCompare but were found via searching the VectorBase server for additional chemosensory genes. In total, 206 chemosensory genes (including all isoforms coding for chemoreceptors) were identified by this process. These were all filtered out of the VectorBase annotated transcripts and GFF files before the [19] chemoreceptors were added, thus eliminating redundancies in the merged GFF file (Supplementary 1). The chemoreceptor coding sequences described by [19] did not include UTR sequences. To ensure uniformity, UTRs were trimmed using a custom python script produced by ChatGPT (Supplementary 2) from all transcripts (Figure 1). *Ae. aegypti* cDNA, 5’UTRs, and 3’UTRs were all extracted using Ensemble Metazoa BioMart. These files were placed into the working directory with the .py script, and a trimmed transcriptome was produced. In addition to generating a new GFF file to include the updated chemoreceptor annotations from [19], a new GAF file (Supplementary 3) and annotated transcripts file (Supplementary 4) were also produced, reflecting the same merge process used in creating the GFF file. Importantly, in the paper by [19] no standardized Gene IDs were given to novel chemosensory genes. Only gene descriptions were provided. Therefore, in the GFF, GAF, and transcriptomics FASTA files, all Vectorbase gene IDs were replaced with gene descriptions in leu of gene IDs. Therefore, all R scripts to utilize these files call chemosensory genes by their gene descriptors and not by their Vectorbase IDs.

The novel chemoreceptor genes from [19] did not have any associated GO terms. GO terms were assigned to chemosensory genes in the GAF file first by providing peptide sequences to the InterProScan tool [54], which automatically assigns GO terms. However, InterProScan did not assign GO terms to all of the new genes. Subsequently Clustal Omega was used to align peptide sequences and produce a phylogenetic tree rooted to alpha tubulin for each chemosensory receptor class (Supplementary 5-7). Genes without GO terms were assigned the same GO terms as the closest neighbor in each phylogenetic tree, to infer GO terms by sequence similarity (Figure 1). The GAF file was used in subsequent TopGO analysis.

#### 2.6.3 RNAseq analysis

Scripts for RNAseq analysis were written in R with the help of ChatGPT as an aid for coding. RNAseq data was not provided to ChatGPT, and the AI system was only used to create and modify scripts, which are all provided in the supplemental files. Code was developed to 1) generate heatmaps of the various chemosensory genes to compare transcript abundance across strains, 2) make Venn Diagrams to summarize significant differences across strains in DESeq2 analysis, 3) make volcano plots with chemosensory genes and Transcription Factors (TFs) highlighted, and 4) perform TopGO analysis of the differentially expressed genes (DEGs) to identify enriched gene ontology terms that were over-expressed or under-expressed in each mutant strain. All R scripts are included in Supplementary 10.

Visualization of the transcript abundance of chemosensory genes was created by plotting the Log10(avgTPM+1) of each transcript. For visualization in the heatmap, genes with lower expression in all strains were filtered to highlight transcripts that may be biologically relevant in the antennal transcriptome. This clearly shows the expression of chemosensory genes in the *Ae. aegypti* antennae and allows for direct comparisons across strains.

Volcano plots were constructed using the results from DESeq2 at the gene level, and extreme log2 Fold Change (absolute value >20) was excluded after they were identified as outliers (high variability within strains, with 0 read count in multiple samples leading to inflated log2 Fold Change values).

### 2.7 Transcription factor analysis

To identify regulatory genes that may coordinate with the co-receptors to promote chemoreceptor expression, we queried the DESeq2 dataset to identify transcription factors (TFs) and chromatin remodelers that were differentially expressed. First, a list of 1078 *Ae. aegypti* transcription factors was obtained from the CIS-BP Database, as well as from a literature search for TFs associated with chemosensory regulation and development. Members from this list were examined to identify TFs that were differentially expressed. The threshold for differential expression was the same as above, with a Log_2_FoldChange > |1| and a pAdj < 0.05.

Two homologs for *D. melanogaster Mip120* were differentially expressed in opposite directions. These included *AAEL020020* and *AAEL005893*. Sequence alignments showed 33 SNPs in the coding regions, and large differences in the intronic regions. Because *AAEL005893* was not mapped to a chromosome, we considered the possibility that there was only one gene in reality, and that *AAEL005893* was simply an artifact found in the unmapped scaffold. To test for the presence of two genes, primers were designed such that one primer annealed to an exonic region with no variability, while a two gene specific primers were designed to map to variable intronic regions. Two such pairs of primers were constructed. Primer sequences were as follows: Mip120_Fwd1 – 5’-CAA ACA ATG GAG GAG CTT GG-3’; AAEL005893_intron_Rev1 – 5’-TTG TCC AAT ACT GTA GGT CC-3’; AAEL020020_intron_Rev1 – 5’-GCT ACG AGG GGA AGT GTA AA-3’; AAEL020020_intron_Fwd2 – 5’-CCT TCC TAG TCA AGT CTT TAT G-3’; AAEL005893_intron_Fwd2 – 5’-CGT CTA ATC TGA TCT GAA TCT C-3’; Mip120_Rev2 –5’-GCT CGA ACG AAA TTT GCT GG-3’. PCR was carried out using the following parameters. 1. Initial denaturation (95⁰C – 2 minutes); 2. Denaturation (95⁰C – 15 seconds); 3. Annealing (56⁰C – 15 seconds); 4. Extension (72⁰C – 20 seconds); 5. Cycle (Steps 2-4 35X total); 5. Final Elongation (72⁰C – 5 minutes). The PCR products were run on a 1% agarose gel and visualized with the Gel Doc EZ Gel Documentation System (BioRad).

Additionally, a Bowtie2 [55] alignment was performed to align RNAseq reads to exon 3 (an exon with a relatively higher abundance of SNPs between the two genes) of *AAEL020020* or *AAEL005893*. Read mapping was visualized in SnapGene to observe proportions of SNPs observed in the reads.

### 2.8 Upstream motif analysis

A python script (Supplementary 11) was written with the help of ChatGPT to read both the *Ae. aegypti* genome .fasta file and the updated .gff3 file (see section 2.6.1), to identify transcription start sites (TSS), and to extract 2,000 bp of DNA sequence upstream (5’) of the TSS. In some cases, less than 2,000 bp would be extracted if the TSS was located near the end of the chromosome. From the DESeq2 analysis (see section 2.6.2), four lists DEGs were generated: *Orco*^−/−^ under-expressed, *Orco*^−/−^ over-expressed, *Ir8a*^−/−^ under-expressed, and *Ir8a*^−/−^ over-expressed. Promoters were filtered to produce four promoter files, one for each list of DEGs. The web-based tool XSTREME (Motif Discovery and Enrichment Analysis) from the MEME Suite 5.5.7 was used for comprehensive motif analysis on each set of DEGs, using default settings [56]. Briefly, XSTREME limited the search to E-values < = 0.05, with a motif width of 6 – 15. The background was created by shuffling input sequences with a Markov order = 2. Motifs meeting the threshold were automatically subjected to Tomtom, the MEME Suite’s tool to compare discovered motifs to known motifs. The set of known motifs was set to JASPAR (non-redundant) DNA: JASPAR CORE (2022): [57]. After obtaining results from the motif analysis, motifs were compared to identify those that were common between each of the four categories of DEGs.

## 3. Results

### 3.1. Strain Confirmation

We analyzed differential gene expression in two mutant strains by comparing their transcriptomes to the Orlando wildtype. The *Orco*^−/−^ strain contains a 16 bp deletion in the first exon, for which there is no read coverage (Figure S1), corresponding to the deletion coordinates [37]. The *Ir8a*^−/−^ strain includes an inserted cassette with the polyubiquitin promoter, *dsRED*, and *SV40* [27]. As expected, transcript coverage increased dramatically, downstream of the polyubiquitin promoter, with coverage spanning the *dsRED* sequence and 3’ exons (Figure S2). We further confirmed the identity of the *Ir8a*^−/−^ strain by examining larvae under a fluorescent microscope (excitation 558 nm, emission 583 nm) to visualize the *dsRED* expression (Figure S3). Of note, *dsRED* is observable under visible light in a dissection microscope, allowing for simplified verification of the strain while dissecting antennae.

### 3.2 Sample Statistics

We performed antennal RNAseq profiling on mated, non-blood-fed female mosquitoes that were aged 5-7 DPE. Our analysis compared the antennal transcriptome of *Orco*^−/−^ and *Ir8a*^−/−^ to the Orlando wildtype. After performing RNAseq for each strain (at least four replicates of each strain), the libraries were subjected to Principal Component Analysis within the DESeq2 platform. The PCA plot revealed a clear separation between the Orlando strain and the two mutant strains, with less separation between the two mutant strains (Figure S4).

### 3.3 Odorant receptors are under-expressed in Aedes aegypti Orco^−/−^

Heat maps of the TPM values for the major chemoreceptor families were generated to visualize gene expression comparisons across samples (Figure 2). Potential differences in magnitude in wild type versus mutant antennae across all three gene families were broadly evident, even before differential expression analysis was performed (Figure 2). DESeq2 comparisons revealed significant differential expression of genes between strains (Supplementary 13,14). Genes were classified as differentially expressed if p-adj < 0.05 and absolute log2-fold-change was >1 [58]. More than 1700 DEGs were identified across samples (Figure 3). Chemoreceptor genes were significantly underrepresented in the *Orco*^−/−^ strain, supporting our initial hypothesis (Figure 3, 4). Fifty-one Ors, including *Orco*, 18 Irs, including *Ir8a*, and 5 Grs were significantly under-represented, while *Or36*, *Or42*, *Or72*, and *Or125* were significantly over-represented (Figure 3, 4).

**Figure 2.**
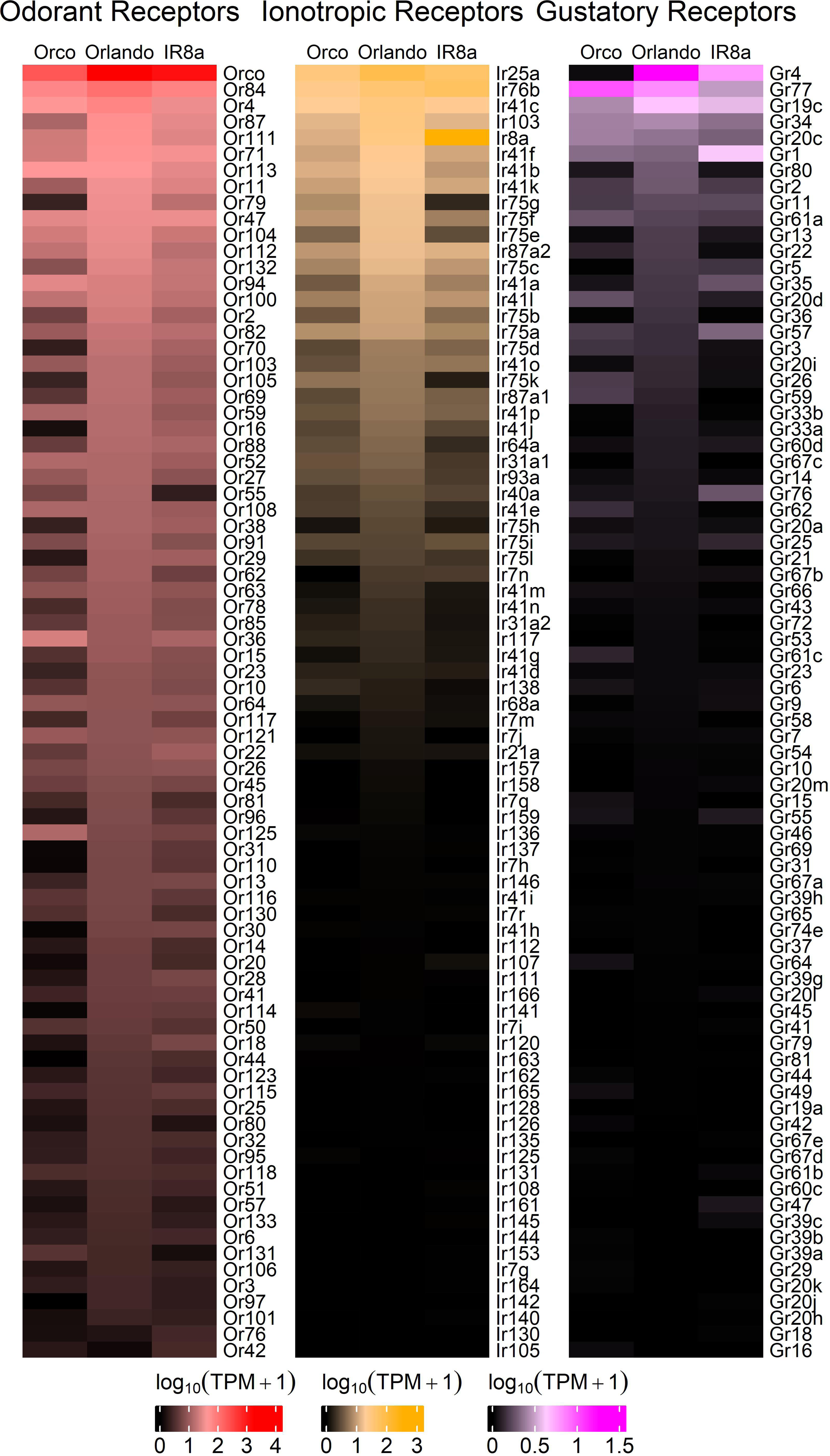
Heatmaps for each chemosensory gene family. Transcript abundances were compared across strains, showing that tuning Ors are broadly under-expressed in *Orco*^−/−^, followed by tuning Irs. In the *Ir8a*^−/−^ mutant strain, tuning Irs are generally under-expressed, followed by tuning Ors. The log10(TPM+1) of each transcript was plotted. Here, the top eighty transcripts are represented, with transcripts that are not highly expressed in all three strains filtered out. Expanded heatmaps are provided in Figure S7.

**Figure 3.**
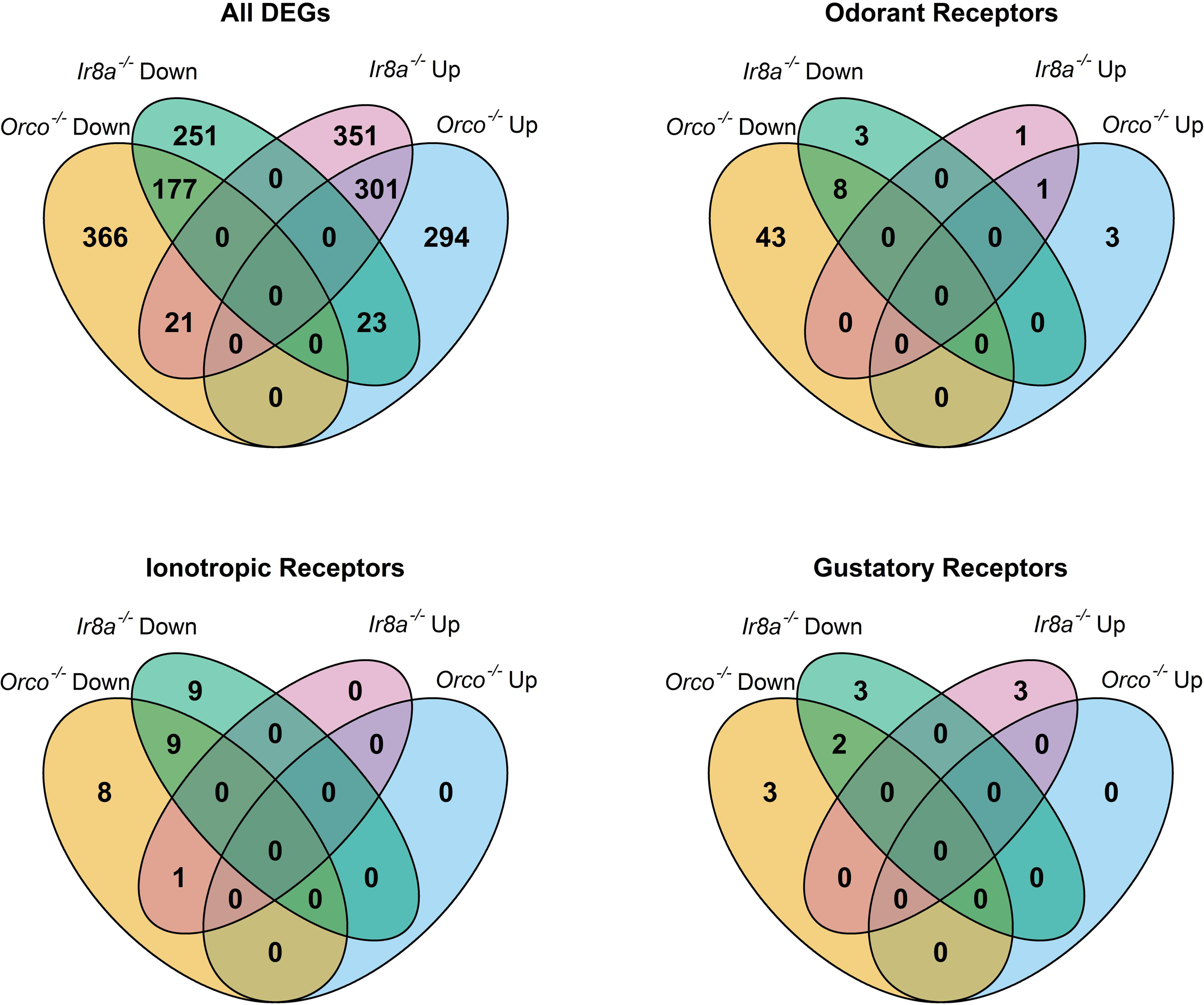
Venn diagrams showing overlap of DEGs across mutant antennae samples as identified by DESeq2 analysis. In each diagram, the regions are labeled by the mutant strain and whether the genes are significantly over-represented (Up) or under-represented (Down) when compared to wild type. All DEGs are considered, as well as subsets of chemoreceptor genes including Ors, Irs, and Grs.

**Figure 4.**
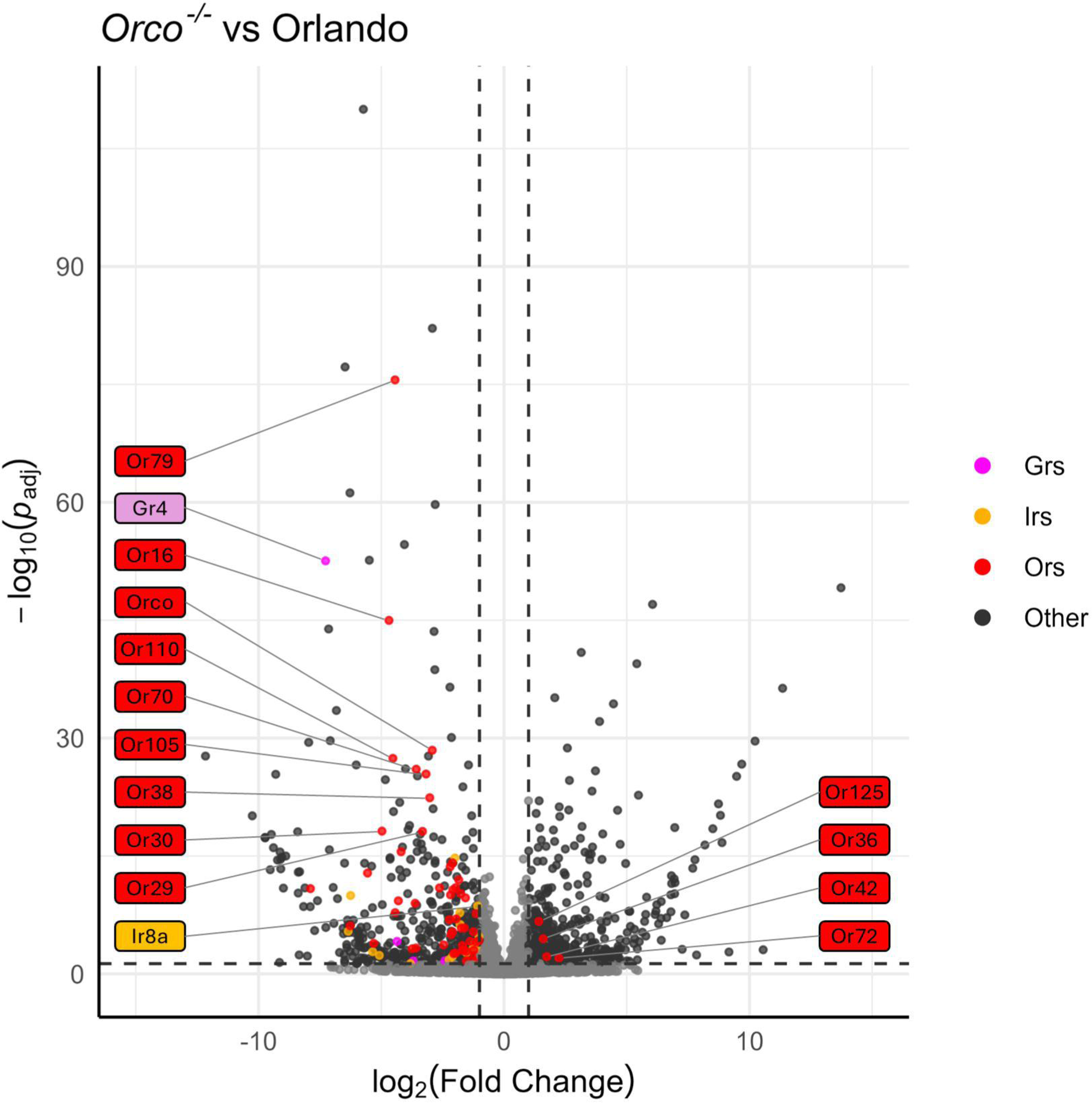
Volcano plot of DESeq2 analysis, comparing Orlando wild type with *Orco*^−/−^. Tuning chemoreceptors, specifically OrXs, are broadly under-represented in *Orco*^−/−^. Odorant receptors are highlighted in red, ionotropic receptors are highlighted in gold, and gustatory receptors are highlighted in magenta. Dotted horizontal and vertical lines represent the thresholds for consideration as DEGs (padj <0.05, absolute log_2_(Fold Change) >1). The 10 chemoreceptors with the lowest padj values are labeled, as well as any chemoreceptor that is significantly over-represented. Co-receptors *Orco* and *Ir8a* are labeled regardless of significance. Labels are colored according to chemoreceptor class.

A TopGO analysis (Supplementary 15,16) was performed to identify the Molecular Functions of genes that were significantly under-or over-represented in the *Orco*^−/−^ strain. Strikingly, among the under-represented GO terms, 17 of 25 with the lowest p-values were descriptive terms for chemoreceptors. These included terms such as “olfactory receptor activity” (p = 1.71E-42), “odorant binding” (p = 2.58E-31), “ligand-gated channel activity” (p = 3.92E-08), and “glutamate receptor activity” (p = 3.78E-4) (Figure 5). This analysis shows that not only are the molecular functions of olfaction significantly reduced in *Orco* mutants, but that only a few other molecular functions are affected to the same level of significance (p<0.005). GO terms associated with over-represented DEGs in *Orco*^−/−^ antennae were associated with peptidases (serine-type endopeptidase activity: p = 1.23e-6), suggesting that in *Orco* mutant antennae, proteolytic cleavage is increased (Figure S5).

**Figure 5.**
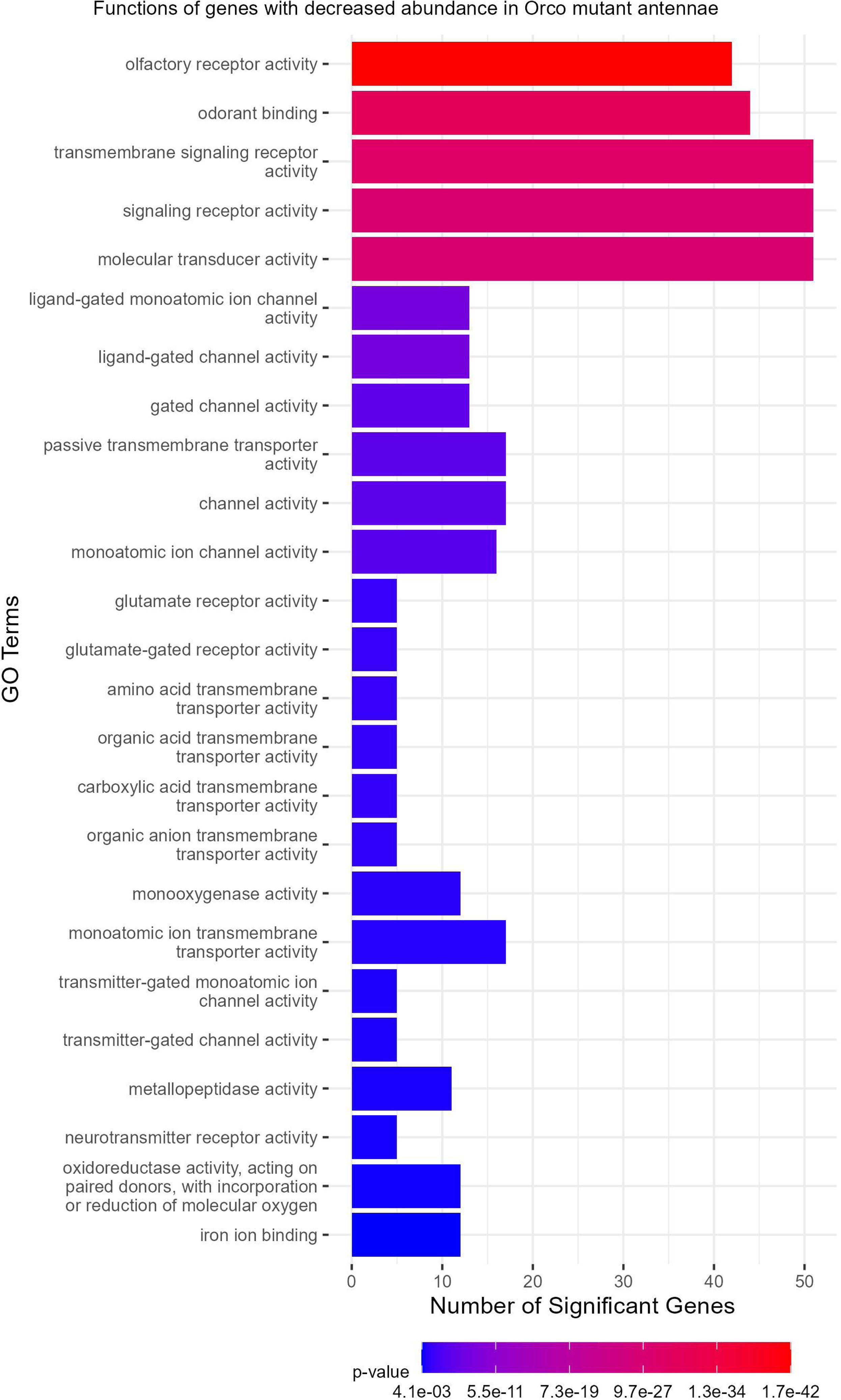
TopGO analysis of DEGs. Olfactory receptor functions are representative of the under-expressed genes in the Orco mutant antennae. TopGO analysis was performed at the Molecular Functions level, and the updated GAF file (including the most recently updated chemoreceptor genes) was used to map GO terms to genes. All GO terms with a classic Fisher’s p-value <0.005 were plotted. The color map from red to blue represents the p-value for the enrichment of each term, with the lowest p-values colored in red.

### 3.4 Chemoreceptor dysregulation in Aedes aegypti Ir8a^−/−^

Similar to *Orco^−/−^*, under-expression of chemosensory genes and over-expression of peptidases was observed in the *Ir8a^−/−^* antennae (Figures 6 and S6). For example, 11 Ors (excluding *Orco*), 18 Irs, and 5 Grs were under-represented (Figures 3 and 6). Conversely, *Or42*, *Or76*, *Ir8a*, *Gr1*, *Gr57*, and *Gr76* were over-represented in the *Ir8a*^−/−^ strain (Figures 3 and 6).

**Figure 6.**
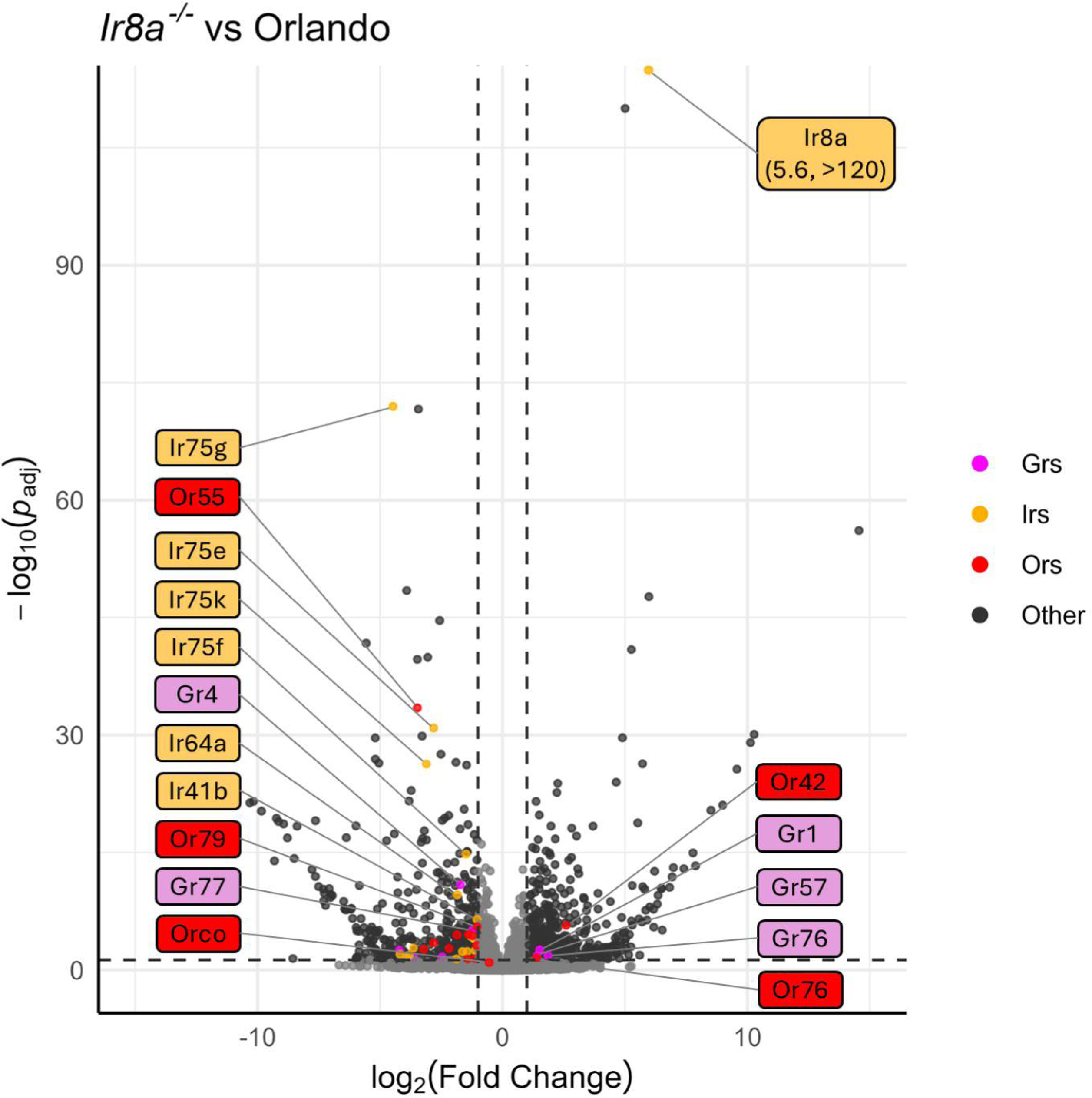
Volcano plot of DESeq2 analysis, comparing Orlando wild type with *Ir8a*^−/−^. Tuning chemoreceptors, specifically IrXs, are broadly under-represented in *Ir8a* mutants. Odorant receptors are highlighted in red, ionotropic receptors are highlighted in gold, and gustatory receptors are highlighted in magenta. Dotted horizontal and vertical lines represent the thresholds for consideration as DEGs (padj <0.05, absolute log_2_(Fold Change) >1). The 10 chemoreceptors with the lowest padj values are labeled, as well as any chemoreceptor that is significantly over-represented. Co-receptors *Orco* and *Ir8a* are labeled regardless of significance. Labels are colored according to chemoreceptor class.

Interestingly, in the *Ir8a*^−/−^ strain the *Ir8a* gene was significantly over-expressed compared to the wildtype gene in Orlando (Figure 5). This overexpression of *Ir8a* is explained by the presence of the polyubiquitin promoter in the *dsRED* cassette, which drives the expression of downstream genes in all tissues. This promoter thus drives the expression of *dsRED*, as well as the downstream out-of-frame *Ir8a* exons (Figure S3).

As in the *Orco*^−/−^ strain, the TopGO analysis (Supplementary 17,18) identified peptidase activity as a functional description of genes with increased abundance in *Ir8a*^−/−^ antennae (Figure S6). In the over-represented *Ir8a*^−/−^ DEGs, Serine-type endopeptidase activity was the most significant GO term (p = 4.18E-14; Figure 7). In the under-represented DEGs, GO terms descriptive of chemoreceptors and ionotropic receptors were identified (Figure 6). Ligand-gated Monoatomic Ion Channel Activity was the most significantly enriched GO term in the under-expressed DEGs (p = 1E-6; Figure 7).

**Figure 7.**
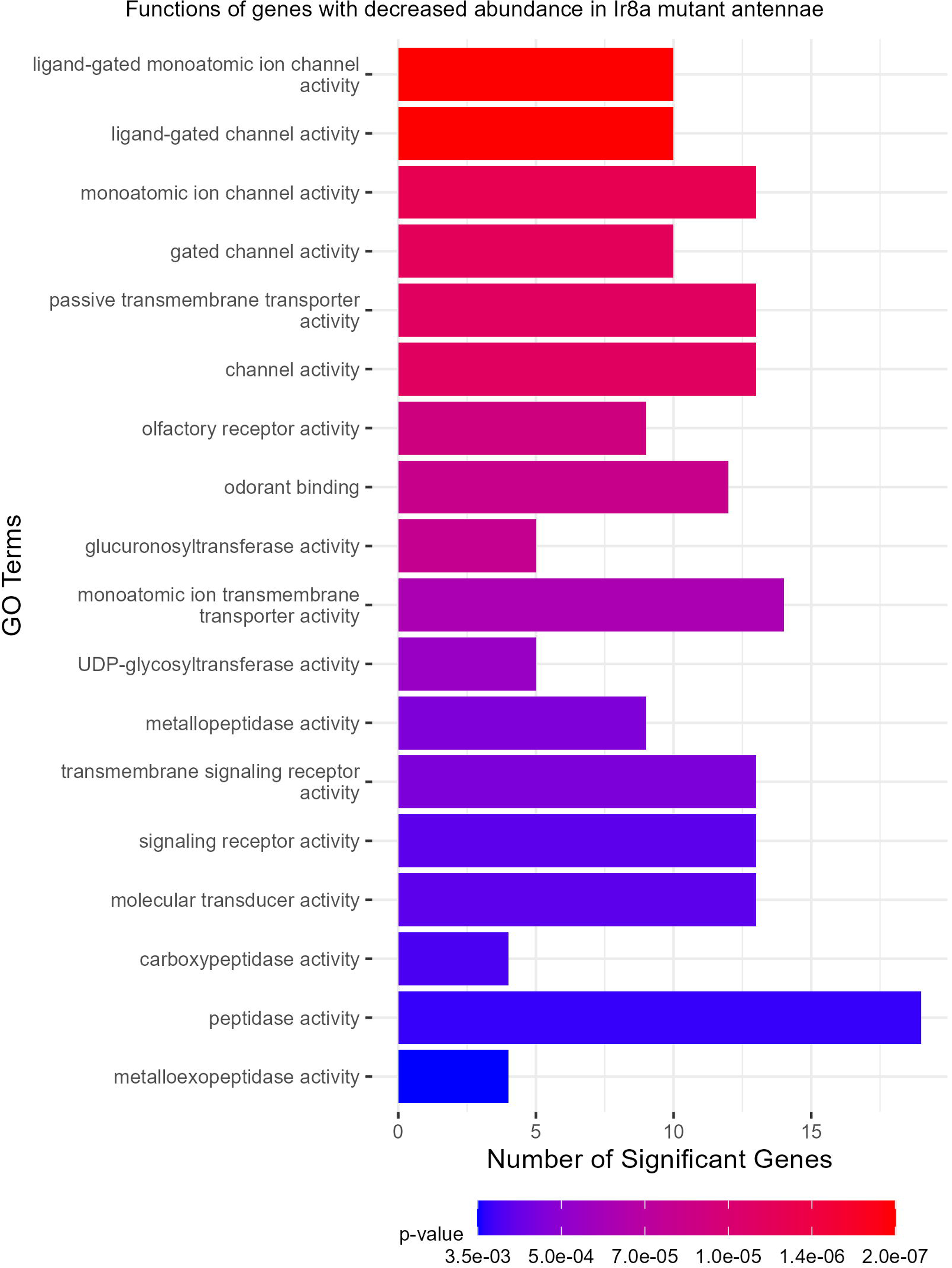
TopGO analysis of DEGs. Ligand-gated ion channel functions are representative of the under-expressed genes in the Ir8a mutant antennae. TopGO analysis was performed at the Molecular Functions level, and the updated GAF file (including the most recently updated chemoreceptor genes) was used to map GO terms to genes. All GO terms with a classic Fisher’s p-value <0.005 were plotted. The color map from red to blue represents the p-value for the enrichment of each term, with the lowest p-values colored in red.

### 3.5 Molecular pathways and transcription factors involved in chemosensory regulation

To identify molecular pathways involved in chemosensory regulation, various transcription factors (TFs) and chromatin remodeling genes were queried for differential expression from the RNAseq dataset. Genes from the *MMB/dREAM* complex, the *miR-279* pathway, as well as various TFs were differentially expressed (Figure 8, Table 1).

**Figure 8.**
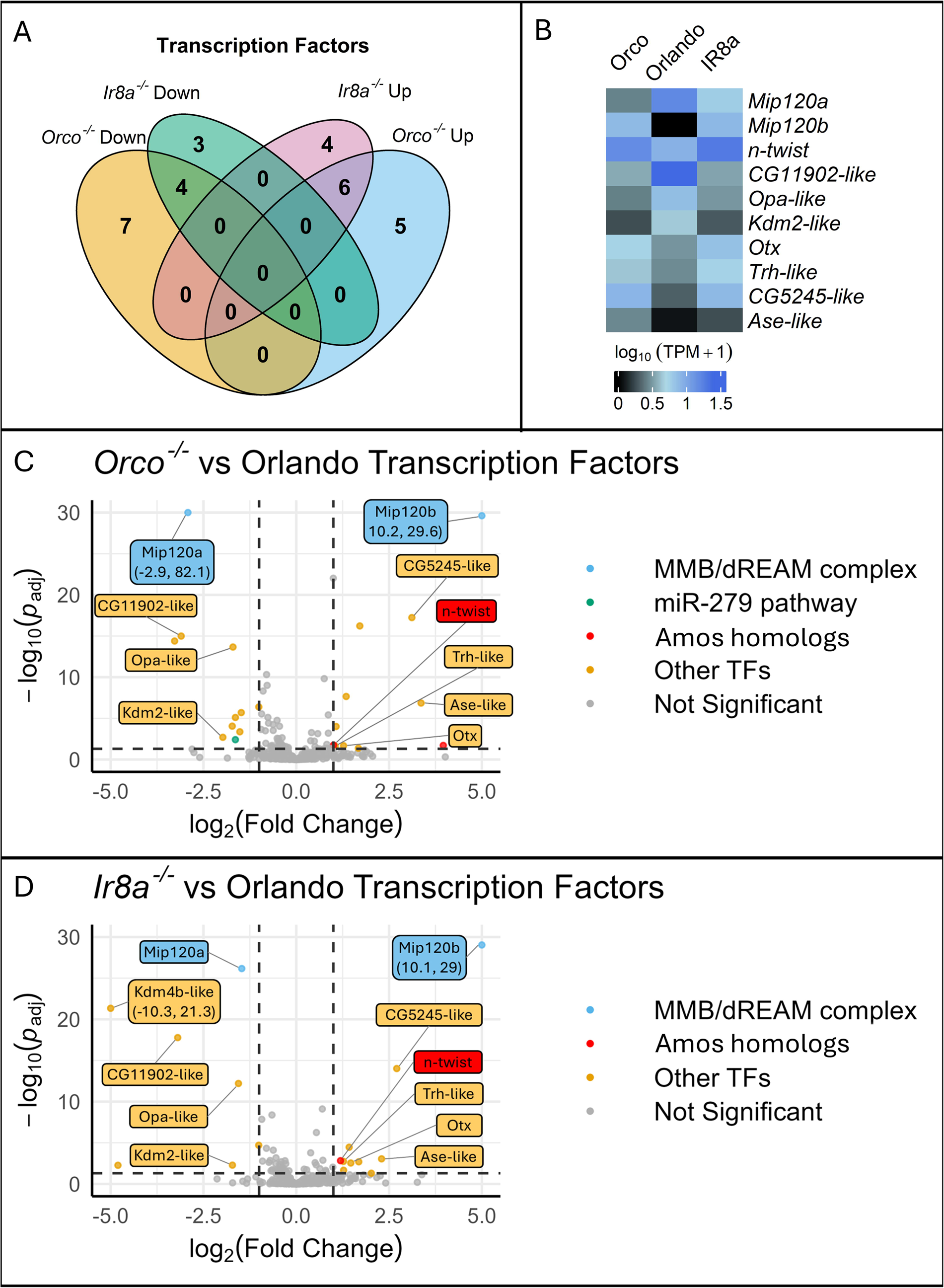
Visualization of the transcription factor differential expression shows two Mip120 homologs (Mip120a and Mip120b), members of the MMB/dREAM complex, along with other TFs including an n-twist, a neuronal differentiation TF with sequence similarity to the D. melanogaster chemosensory TF amos to be significantly differentially expressed. A. A Venn Diagram shows the overlap between Orco−/− and Ir8a−/− mutants of differentially expressed TFs compared to the wildtype. B. A heatmap of the relative expression levels for the 10 differentially expressed genes in common to both co-receptor mutants. C-D. Volcano plots showing differentially expressed TFs in Orco−/− or Ir8a−/− relative to the wildtype. Any datapoints that were not TFs were filtered out prior to visualization. Labels were added to the ten DEG TFs in common to both mutant strains. The Y-axis was confined to 30, and the X-axis was confined to |5.0|. Any points with X or Y values outside of those limits were plotted on the edge of the graph, labels were added, and the X,Y coordinates were included in the label.

**Table 1.**
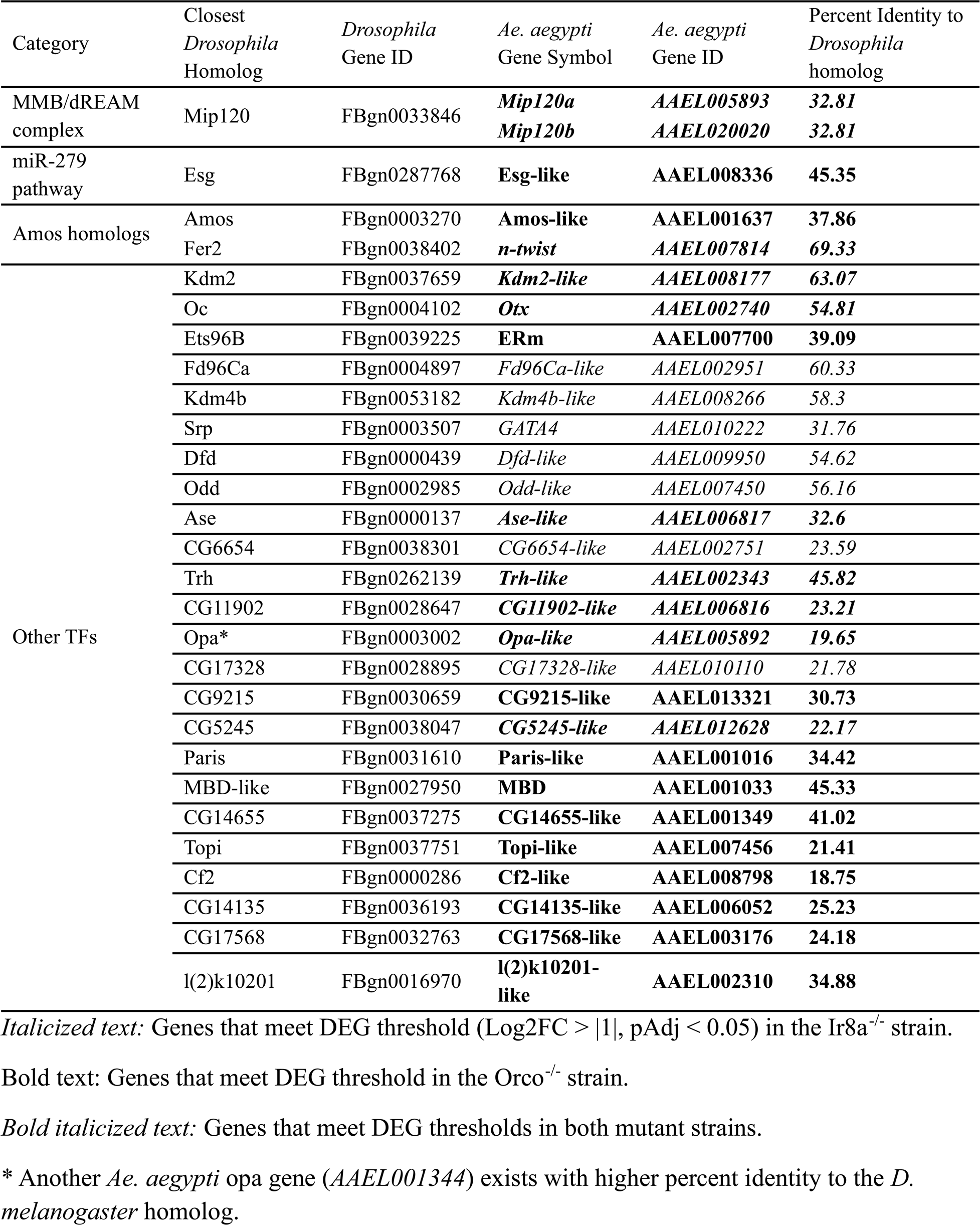
Transcription factors and chromatin remodeling genes implicated in chemoreceptor regulation in *Aedes aegypti* mosquitoes.

| Category | Closest<br>*Drosophila*<br>Homolog | *Drosophila*<br>Gene ID | *Ae. aegypti*<br>Gene Symbol | *Ae. aegypti*<br>Gene ID | Percent Identity to<br>*Drosophila*<br>homolog |
| --- | --- | --- | --- | --- | --- |
| MMB/dREAM<br>complex | Mip120 | FBgn0033846 | *Mip120a* | **AAEL005893** | **32.81** |
|  |  |  | *Mip120b* | **AAEL020020** | **32.81** |
| miR-279<br>pathway | Esg | FBgn0287768 | **Esg-like** | **AAEL008336** | **45.35** |
| Amos homologs | Amos | FBgn0003270 | **Amos-like** | **AAEL001637** | **37.86** |
|  | Fer2 | FBgn0038402 | ***n-twist*** | **AAEL007814** | **69.33** |
| Other TFs | Kdm2 | FBgn0037659 | ***Kdm2-like*** | **AAEL008177** | **63.07** |
|  | Oc | FBgn0004102 | ***Otx*** | **AAEL002740** | **54.81** |
|  | Ets96B | FBgn0039225 | **ERm** | **AAEL007700** | **39.09** |
|  | Fd96Ca | FBgn0004897 | *Fd96Ca-like* | AAEL002951 | 60.33 |
|  | Kdm4b | FBgn0053182 | *Kdm4b-like* | AAEL008266 | 58.3 |
|  | Srp | FBgn0003507 | *GATA4* | AAEL010222 | 31.76 |
|  | Dfd | FBgn0000439 | *Dfd-like* | AAEL009950 | 54.62 |
|  | Odd | FBgn0002985 | *Odd-like* | AAEL007450 | 56.16 |
|  | Ase | FBgn0000137 | ***Ase-like*** | **AAEL006817** | **32.6** |
|  | CG6654 | FBgn0038301 | *CG6654-like* | AAEL002751 | 23.59 |
|  | Trh | FBgn0262139 | ***Trh-like*** | **AAEL002343** | **45.82** |
|  | CG11902 | FBgn0028647 | ***CG11902-like*** | **AAEL006816** | **23.21** |
|  | Opa* | FBgn0003002 | ***Opa-like*** | **AAEL005892** | **19.65** |
|  | CG17328 | FBgn0028895 | *CG17328-like* | AAEL010110 | 21.78 |
|  | CG9215 | FBgn0030659 | ***CG9215-like*** | **AAEL013321** | **30.73** |
|  | CG5245 | FBgn0038047 | ***CG5245-like*** | **AAEL012628** | **22.17** |
|  | Paris | FBgn0031610 | **Paris-like** | **AAEL001016** | **34.42** |
|  | MBD-like | FBgn0027950 | **MBD** | **AAEL001033** | **45.33** |
|  | CG14655 | FBgn0037275 | ***CG14655-like*** | **AAEL001349** | **41.02** |
|  | Topi | FBgn0037751 | **Topi-like** | **AAEL007456** | **21.41** |
|  | Cf2 | FBgn0000286 | **Cf2-like** | **AAEL008798** | **18.75** |
|  | CG14135 | FBgn0036193 | ***CG14135-like*** | **AAEL006052** | **25.23** |
|  | CG17568 | FBgn0032763 | ***CG17568-like*** | **AAEL003176** | **24.18** |
|  | l(2)k10201 | FBgn0016970 | **l(2)k10201-like** | **AAEL002310** | **34.88** |

*Italicized text:* Genes that meet DEG threshold ( $\text{Log2FC} > |1|$ , $\text{pAdj} < 0.05$ ) in the Ir8a<sup>-/-</sup> strain.
**Bold text:** Genes that meet DEG threshold in the Orco<sup>-/-</sup> strain.
***Bold italicized text:*** Genes that meet DEG thresholds in both mutant strains.
\* Another *Ae. aegypti* opa gene (AAEL001344) exists with higher percent identity to the *D. melanogaster* homolog.

Most notably, there were two *Ae. aegypti* homologs to the *Drosophila melanogaster Mip120* gene. *Mip120* is part of the *MMB/dREAM* complex and was previously characterized as a repressor of *Gr63a* in inappropriate neurons [59]. The *MMB/dREAM* complex was also implicated as a chemoreceptor regulator for other Ors [59]. *Mip120b* (*AAEL020020*), one of the *Mip120* homologs, was significantly over-expressed (Log_2_FoldChange >10) in both mutant strains (Figure 8, Table 1). An additional *Mip120* homolog, *Mip120a* (*AAEL005893*) was significantly under-represented. *Mip120b* was located on chromosome 3, but *Mip120a* was unmapped to a chromosome and was annotated to the NIGP01000892 scaffold. Therefore, we reasoned that *Mip120a* may not be a real gene, but merely an artifactual copy of *Mip120b*. Therefore, we sought to determine if one or two genes were present in the *Aedes aegypti* genome. The exonic regions of the genes were nearly identical, with only 33 SNPs between the two coding sequences. The annotated introns were different however. We designed two sets of primers. In set 1, a forward primer was designed to anneal to an exonic region shared by both genes. Two separate reverse primers were designed to anneal to gene specific regions in the introns. In set 2, the reverse primer annealed to a shared exonic regions, while the forward primers annealed to gene specific primers. When the genes were PCR amplified, bands were present for all primer sets, confirming the presence of both genes (Figure S8).

We found it remarkable that in Orlando wild-type expression results, *Mip120b* had no read counts that were mapped to the gene, while hundreds of reads were mapped to *Mip120a* (Figure 8B), since the transcript sequences were nearly identical. In the mutant strains, reads mapped to both genes (Figure 8B). With only 33 SNPs found in the coding regions between both genes, we doubted Salmon’s ability to accurately map reads between these ambiguous transcripts. We ran a Bowtie2 alignment of reads to either *Mip120a* or *Mip120b* exon 3 (a region with relatively more SNPs). We then visualized the read alignment in SnapGene. In Orlando wild-type, all of the reads that mapped to *Mip120a/b* only contained *Mip120b* specific SNPs. No *Mip120a* polymorphisms were observed in Orlando transcripts. When examining mutant strains, the reads contained both *Mip120a* and *Mip120b* polymorphisms. This established our findings that, though *Mip120a* and *Mip120b* are similar at the transcript level, Salmon was able to accurately map reads to each transcript. This verifies that both genes exist in the genome and that they are differentially expressed in opposite directions in both mutant strains. In *Ae. aegypti,* these genes had not yet been given a name. We therefore named the *AAEL005893* and *AAEL020020* genes *Mip120a* and *Mip120b,* respectively.

Additionally, an *Esg* homolog was significantly under-represented in *Orco*^−/−^ mutants, implicating the *miR-279* pathway in chemoreceptor regulation (Figure 8, Table 1). Two *Amos* homologs, *amos-like* (*AAEL001637*) and *n-twist* (*AAEL007814*), were significantly over-represented in Orco^−/−^ alone and in both mutants, respectively (Figure 8, Table 1). *Kdm2* and *Oc* homologs were identified as DEGs in both mutant strains and were under-or over-represented, respectively (Table 1). *Ets96B* was significantly over-represented in *Orco*^−/−^ mutants, and TFs *Fd96Ca*, *Kdm4b*, *Srp*, *Dfd*, and *Odd* were identified as DEGs in *Ir8a*^−/−^ mutants (Table 1).

### 3.6 Molecular pathways and transcription factors involved in chemosensory regulation

The XSTREME motif discovery and enrichment algorithm was used to identify potential cis-regulatory elements in the regions 2,000 bp upstream of the TSS for DEGs. Within these promoter regions, eight motifs were identified in *Orco*^−/−^ under-expressed DEGs, ten in *Orco*^−/−^ over-expressed DEGs, three in *Ir8a*^−/−^ under-expressed DEGs, and twenty-five in *Ir8a*^−/−^ overexpressed DEGs. Enriched motifs were compared between DEG categories. While the threshold for XSTREME analysis was set at *E<0.005*, the algorithm also returns the top three motifs identified by SEA (Simple Enrichment Analysis), even if the *E* values are greater. Five motifs are highlighted below for further consideration. Two motifs, the first similar to those recognized by *achi* and *vis* in *D. melanogaster* and the second with no known TFs associated, were found in the upstream regions of Orco^−/−^ under-expressed DEGs and Ir8a^−/−^ over-expressed DEGs (Figure 9). Three motifs were enriched in the upstream regions of over-expressed DEGs in both mutants. These motifs are identical to the *D. melanogaster* motifs recognized by *CG4328-RA*, *Clamp*, and *su(Hw)*.

**Figure 9.**
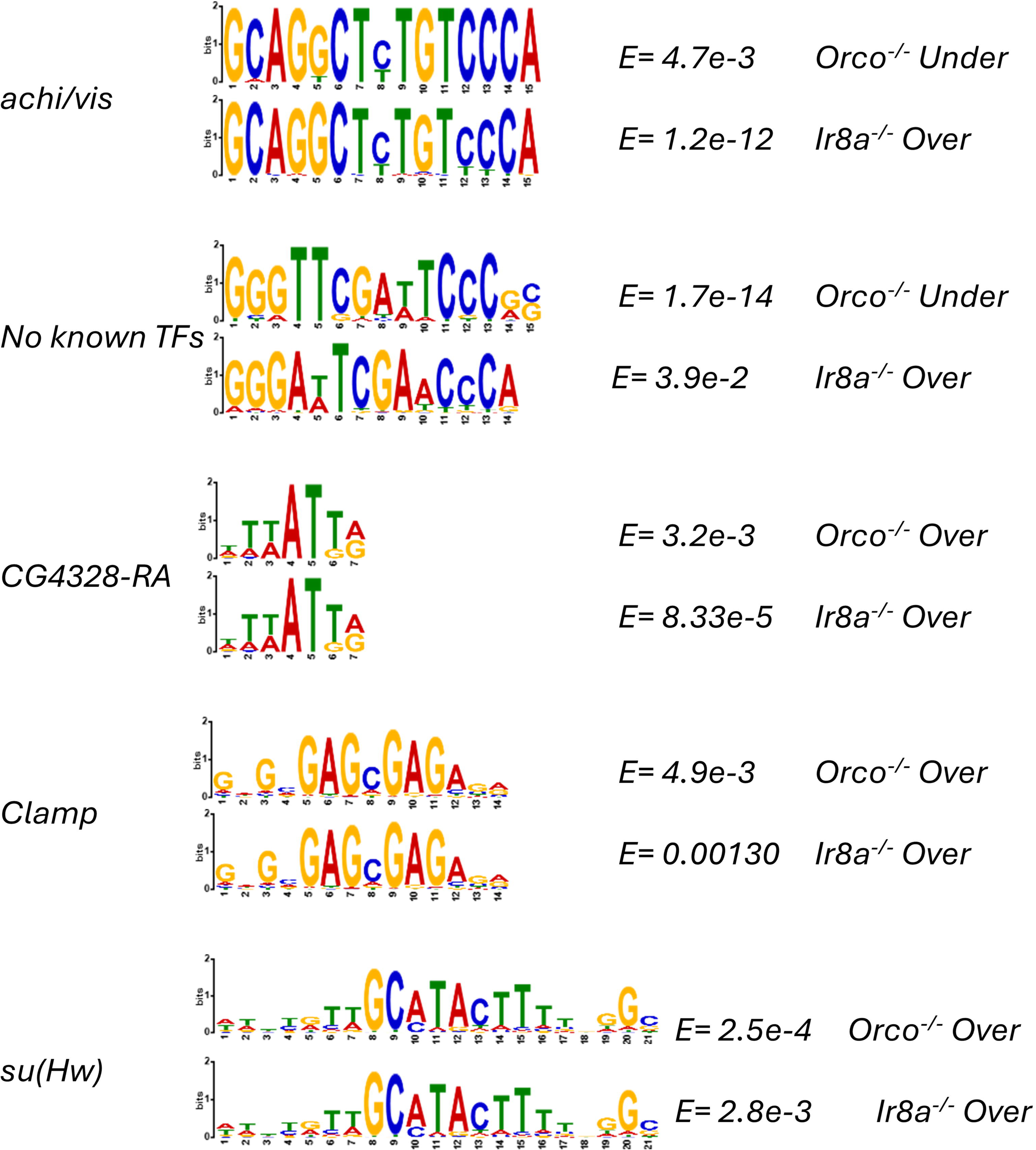
XSTREME motif analysis reveals that similar motifs are enriched in both co-receptor mutants. Motifs that are significantly (*E<0.*05) enriched and are represented in both mutants are displayed. If an insect TF is known to bind to a motif, the name of the TF is printed to the right. The E values from the XSTREME analysis are provided to the right, as well as the set of DEG upstream regions from which the motif was discovered.

## 4. Discussion

In this study, we examined the expression of chemosensory genes in the antennae of female *Ae. aegypti*, comparing a wildtype strain with two co-receptor mutant strains, *Orco*^−/−^ and *Ir8a*^−/−^. Our analysis reveals that tuning receptors are under-expressed in mutant antennae compared to a wildtype strain and provides evidence in support of the hypothesis that functional co-receptors are required for expression of tuning receptors. Two non-exclusive explanatory models may account for these observations. First, the evidence may indicate pleiotropic roles for co-receptors, namely that of coordinating expression of their cognate tuning receptors. Alternatively, the under-representation of tuning receptors could be the result of the degeneration of neurons expressing either *Orco* or *Ir8a* due to physiological inactivity. Here, we contextualize these two models within the body of literature on this subject and propose future directions for research.

In *D. melanogaster*, when *Orco* or *Ir8a* are functionally disrupted, the transport of cognate tuning receptors from the endoplasmic reticulum to the dendritic surface is impaired and expression of the cognate tuning receptors is reduced [36,38,39,44]. Furthermore, when *Ir8a* is knocked out, *Ir64a* protein abundance is reduced in the antennae [38]. When we performed RNAseq differential expression analysis in *Ae. aegypti* antennae, we observed significantly reduced levels of chemoreceptor transcripts. This could indicate that the co-receptors, in addition to directing the transport of tuning receptors, also coordinate their transcription. Such pleiotropy was observed in *D. melanogaster*, where transcript abundance of the cognate coreceptors was reduced, even in neurons that did not undergo neurodegeneration [44]. Our study in *Ae. aegypti* indicates that this phenomenon may not be specific to *D. melanogaster*.

Studies in *D. melanogaster* have also demonstrated that co-receptor expression is required for sensory neuron development or maintenance, as neurodegeneration is observed within the OSNs in early adulthood [43]. In contrast, a large population of OSNs survive, even when the co-receptor is abolished [44]. In *Ae. aegypti* OSN neurodegeneration has not been observed, though this has not been extensively studied [43]. Follow-up studies should evaluate the extent and timing of potential neurodegeneration in *Orco*^−/−^ and *Ir8a*^−/−^ mutant mosquitoes. To do this, the *T2A-QF2* in-frame fusions for *Orco* and *Ir8a* could be utilized [60]. Though these insertions are in-frame, the authors posited that producing a homozygous driver line could result in a loss-of-function phenotype. If such a phenotype is produced, then the homozygous in-frame fusion would be sufficient to track neuronal health. Alternatively, a homozygous driver line for a *T2A-QF2* out-of-frame fusion could be created. This would lead to functional knockouts for each co-receptor gene, while enabling visualization of frameshifted transcript expression in ORNs and IRNs as proxies for neuronal health.

Both models provide explanations for the reduced transcript abundance observed in co-receptor knockout mosquitoes and are supported by findings in *D. melanogaster* [38,39,43,44]. We propose that reduced transcript abundance can be explained by a combination of both models. In such a scenario, a subset of OSNs undergo cell death because of inactivity. Those that survive display a decreased abundance of the cognate transcripts for each co-receptor due to loss of the co-receptor’s coordination of transcription.

We further investigated whether any TFs or chromatin remodeling genes were differentially expressed. We identified a total of 29 regulatory genes that were differentially transcribed. Of these, several TFs had previously been associated with regulation of chemosensory genes, including *amos* homologs, members of the *MMB/dREAM* complex, and a member of the *mrR-279* pathway.[59, 61, 62-].

In *D. melanogaster*, *amos* and *atonal* are two basic-helix-loop-helix transcription factor that have strong similarity to each other [63]. *Amos* is involved in the development of olfactory sensilla in *Drosophila* and together with *atonal* is responsible for the formation of the ORNs and the antennal lobe [64]. Two *amos* homologs were differentially expressed in the mutant strains of mosquitoes. First, *amos-like* (*AAEL001637*), which shares approximately 38 percent identity with *D. melanogaster amos*, is significantly over-represented in *Orco*^−/−^ mutants. Next, *n-twist* (*AAEL007814*), which shares approximately 35 percent identity with *D. melanogaster amos*, is significantly over-represented in both *Orco*^−/−^ and *Ir8a*^−/−^ mutants. *N-twist* is a transcriptional inhibitor, has been shown to affect differentiation of neurons, and shows sequence similarity to *amos* [65]. Whether it directly regulates chemosensory receptors has not been determined. Over-expression of the *n-twist* gene could be due to a feedback loop; when regulatory feedback loops detect under-expression of chemoreceptors, they may increase expression of *amos* homologs in an attempt to increase downstream chemoreceptor expression.

The *microRNA-279* pathway has also been implicated in chemosensory gene regulation, where the miRNA works in conjunction with *Nerfin-1*, *Pros*, and *Esg* in *D. melanogaster* to inhibit the development of CO_2_ ORNs in the maxillary palp [66]. The expression levels of an *Esg* homolog were differentially expressed at lower levels in the *Orco*^−/−^ mutants. Under-expression of this gene does not easily explain the broad under-expression of chemoreceptors in both mutants, but it does implicate a role for this TF in the regulation of ORNs. As posited for the *amos* homolog, it may be possible that when the mosquito regulatory systems detect broad under-expression of chemosensory genes, they repress the *miR-279* pathway to promote expression of receptors and formation of ORNs.

When examining differential expression of regulatory genes, the over-representation of *Mip120b* (*AAEL020020*) in both mutant strains may offer the most straightforward explanation for our observation of chemoreceptor under-expression. *Mip120b* is a homolog of *Mip120*, which is a part of the *MMB/dREAM* complex in *D. melanogaster* [59, 67]. Multiple proteins, including TFs and chromatin remodelers, come together to form the *MMB/dREAM* complex [59, 67]. This complex is composed of *Lin-52*, *Rpd3*, *L(3)mbt*, *Myb*, *E2F2*, *DP*, *RBF1/2*, *Mip40*, *Mip120*, *Mip130*, and *P55/Caf1* [59, 67]. In *D. melanogaster*, *Myb* is necessary for *Gr21a* and *Gr63a* expression, while *Mip120* represses aberrant expression in other neurons [59]. There are two *Mip120* homologs in *Ae. aegypti*: *Mip120a* (*AAEL005893*) and *Mip120b (AAEL020020*).

Interestingly, *Mip120a* is significantly under-represented, while *Mip120b* is significantly over-represented in both mutants. If both genes are involved in repressing the expression of chemoreceptors in inappropriate neurons, then over-expression would lead to lower transcription of chemoreceptors, while under-expression would have the opposite effect. While *Mip120a* is significantly under-expressed relative to wildtype, *Mip120b* is over-represented with a positive fold change that is orders of magnitude larger than the negative fold change of *Mip120a*. In both mutants, the total expression of *Mip120b* is greater than that of *Mip120a.* If *Mip120b* shares the *D. melanogaster Mip120* role as chemoreceptor repressor, then its elevated expression relative to the wildtype could explain the broad under-representation of chemoreceptor genes found in *Orco*^−/−^ and *Ir8a*^−/−^ mutants.

After analyzing TFs that were differentially expressed, promoter regions (defined as 2,000 bp upstream of the TSS) were extracted for over– and under-represented DEGs for both *Orco*^−/−^ and *Ir8a*^−/−^ strains. These were subjected to motif discovery and enrichment analysis using the web-based tool XSTREME, part of the MEME Suite 5.5.7. Comparisons were made between motifs enriched in each different list of DEGs. While it is not possible to draw a clear connection between the motif analysis and the TFs that were differentially expressed, some interesting connections were found between motifs enriched in different categories of DEGs. The comparison of motifs between the *Orco*^−/−^ and *Ir8a*^−/−^ over-expressed genes showed the greatest number of similar motifs, with a total of 9 comparisons made between motifs in each category. Interestingly, the *GATAd-* and *su(Hw)*-like binding sites were the most frequently repeated motifs observed. Furthermore, a striking similarity was observed between the *D. melanogaster E2F2* motif and the *su(Hw)* motif identified in Ir8a^−/−^ over-represented DEG promoters [67].

Recently, a feedback loop was identified in the antennae of *Anopheles gambiae*. When *AgamOr2* was expressed ectopically in all ORNs, the expression of any other Or was significantly repressed [68]. The pattern of under-expressed *Ors* and I*rs* reflected the pattern of under-expressed chemoreceptors in our study [68]. From these data, we conclude that the expression of *Orco* or *Ir8a* is necessary to allow expression of cognate tuning receptors, and that when one tuning receptor is effectively expressed, other tuning receptors are repressed. We propose a model whereby *Orco*, *Ir8a*, and the appropriate tuning receptor for a given OSN coordinate with the *MMB/dREAM* regulatory complex to achieve fine tuning of chemosensory control.

This work expands our understanding of the roles of *Orco* and *Ir8a* co-receptors in insects. Prior work has established the roles of *Orco* and *Ir8a* as co-receptors and has implicated them in the coordinated transport of tuning receptors and maintaining the health of OSNs [38,39,43,44]. Our study expands this pleiotropy to *Ae. aegypti* and suggests TFs and putative TF binding motifs that may play a role in chemoreceptor transcriptional regulation. Whether co-receptor coordination of tuning receptor transcription is limited to dipterans, or if is a general phenomenon across other insect orders remains to be investigated. We also note that chemical response phenotypes associated with the *Ae. aegypti Orco^−/−^* and *Ir8a^−/−^* mutants should be reassessed considering the dysregulation of chemoreceptor genes across families [37, 27, 35].

For example, the mutation of either co-receptor led to decreased abundance of *Gr4*, a homolog of *Drosophila melanogaster Gr64f*, the sugar taste receptor [69, 70]. It is interesting that this receptor would be the most highly expressed GrX in wild-type antennae, and that mutation of co-receptors *Orco* and *Ir8a* would be associated with decreased transcription. The complex interplay of co-receptor and tuning receptor expression suggests caution when interpreting results that might otherwise be assumed to affect only one gene family.

Mosquitoes transmit pathogens that disable or kill millions of people around the world annually [8]. In the ongoing struggle to combat vector-borne diseases, it will be imperative to conceptualize novel means to control vector populations to reduce the public health burden imposed by them. Importantly, such strategies should be designed with the problem of insecticide resistance in mind [71]. The current study suggests that in *Ae. aegypti* there exists a tightly regulated transcriptional control of chemosensory receptors. The data further suggests that *Orco* and *Ir8a* co-receptors are integrally involved in this transcriptional control. Continuing to discover the molecular processes, promoter sequences, transcription factors, and other genes involved in such regulation is of high importance. Elucidation of such molecular pathways could lead to the discovery of novel molecular targets for next-generation pesticides. Insects will often develop resistance to pesticides by accumulating mutations in the target of a pesticide [72]. If there is a high fitness cost for mutation of a particular target, we would predict that selection would occur at a slower rate. Conceptually, next-generation insecticides/repellents could be developed to target one or more of the regulatory processes described in this study and integrated into current intervention strategies to mitigate the effects of resistance. However, if mosquitoes developed resistance via mutations in chemoreceptors or their transcriptional regulators, this might lead to a reduction in their ability to efficiently locate blood hosts. Both outcomes would theoretically decrease the vectorial capacity, either by decreasing populations directly or by decreasing the biting rate [8].

## 5. Conclusions

Our differential expression analysis highlights the aberrant expression profiles of chemoreceptor transcripts in *Ae. aegypti Orco*^−/−^ or *Ir8a*^−/−^ co-receptor mutants. We speculated that mutations in the *Orco* and *Ir8a* co-receptors would lead to the broad dysregulation of tuning Ors and Irs, respectively. Our findings support this hypothesis and suggest a pleiotropic role for olfactory co-receptors in *Ae. aegypti* in coordinating chemosensory transcript expression.

Future studies should expand the transcriptomic analysis and explore mechanisms for the coordination of chemosensory transcript expression. To expand the RNAseq analysis, follow-up experimentation should focus on the *Ir8a*^attP/attP^ strain, as the global expression of the *dsRED* protein may affect the expression of other transcripts. Additionally, the transcriptomes of *Orco*^−/−^ and *Ir8a*^−/−^ mutants should be analyzed across chemosensory tissues, sexes, and under different physiological conditions. Such comprehensive analyses may reveal other novel roles for co-receptors. Moreover, the role of conserved motifs in the promoter regions should be evaluated by performing reporter assays in wildtype and mutant co-receptor strains. Such an analysis would contribute towards a mechanistic understanding of the roles of co-receptors in the coordination of chemoreceptor transcript expression. Finally, the molecular pathway for the regulation of chemoreceptor transcription should be fully elucidated to provide a basis for potential targets of next-generation insecticides and repellents.

## Supplementary Materials

Supplementary 01: Merged_Annotations_AGWG.gff; Supplementary 02: remove_utrs.py; Supplementary 03: Merged_Gene_Ontology_Terms.gaf; Supplementary 04: Merged_Aedes_aegypti_transcriptome.fasta; Supplementary 05: Ors Phylogeny.svg; Supplementary 06: Irs Phylogeny.svg; Supplementary 07: Grs Phylogeny.svg; Supplementary 08: Orco_16_transcriptome.fasta; Supplementary 09: Ir8a_dsRED_Transcriptome.fasta; Supplementary 10: R_scripts.txt; Supplementary 11: extract_upstream_sequences.py; Supplementary 12: Figures S1 – S7.docx; Supplementary 13: DESeq2_results_orco_vs_orlando.csv; Supplementary 14: DESeq2_results_ir8a_vs_orlando.csv; Supplementary 15: Orco_DN_GO_Enrichment_Summary.csv; Supplementary 16: Orco_UP_GO_Enrichment_Summary.csv; Supplementary 17: Ir8a_DN_GO_Enrichment_Summary.csv; Supplementary 18: Ir8a_UP_GO_Enrichment_Summary

## Author Contributions

Conceptualization, Matthew Cooke and R. Jason Pitts; Data curation, Matthew Cooke and R. Jason Pitts; Formal analysis, Matthew Cooke; Funding acquisition, R. Jason Pitts; Investigation, Matthew Cooke and Michael Chembars; Methodology, Matthew Cooke, Michael Chembars and R. Jason Pitts; Project administration, R. Jason Pitts; Resources, R. Jason Pitts; Software, Matthew Cooke; Supervision, R. Jason Pitts; Validation, Matthew Cooke and R. Jason Pitts; Visualization, Matthew Cooke; Writing – original draft, Matthew Cooke; Writing – review & editing, Matthew Cooke, Michael Chembars and R. Jason Pitts.

## Data Availability Statement

Supporting data and results are provided as supplementary materials, as described above. RNAseq reads have been deposited in the National Center for Biotechnology Information, Sequence Read Archive (project PRJNA1249520; sample files SRR33089554-SRR3308569).

## Supporting information

Supplementary XSTREME Motif Analysis

## Acknowledgments

The following reagent was obtained through BEI Resources, NIAID, NIH: *Aedes aegypti* Orlando orco^16^, NR-44378. The Ir8a^dsRED/dsRED^ and Ir8a^attP/attP^ strains were provided by Dr. Matthew DeGennaro (Florida International University). We also thank John Boyi and Everest Castaneda for helpful discussions on bioinformatics analysis and Isuru Gunarathna for assistance in creating a merged gff file for *Ae. aegypti* genome annotations. We acknowledge the use of ChatGPT generative AI to aid in writing R and python scripts.

## Abbreviations

The following abbreviations are used in this manuscript:

Orco: Odorant Receptor co-receptor
OrX: Odorant Receptor Tuning Receptor
IrX: Ionotropic Receptor Tuning Receptor
GrX: Gustatory Receptor Tuning Receptor
DPE: Days Post Eclosion
ZT: Zeitgeber Time
GAF: General Annotation File
GFF: Generic Feature File
OSN: Olfactory Sensory Neuron
ORN: Odorant Receptor Neuron
DEG: Differentially Expressed Gene

