## Supplementary Files 1-18 for "The co-receptors *Orco* and *Ir8a* are required for coordinated expression of chemosensory genes in the antennae of the yellow fever mosquito, *Aedes aegypti*": Supplementary 02□ remove_utrs.py

```
from Bio import SeqIO

def load_fasta_as_dict(fasta_file):
    """Load a FASTA file and return a dictionary with headers as keys and sequences as values."""
    sequences = {}
    for record in SeqIO.parse(fasta_file, "fasta"):
        sequences[record.id] = str(record.seq)
    return sequences

def remove_utrs(cdna_dict, utr5_dict, utr3_dict):
    """Trim 5' and 3' UTR sequences from cDNA sequences."""
    trimmed_sequences = {}
    for header, cdna_seq in cdna_dict.items():
        utr5_seq = utr5_dict.get(header, "")
        utr3_seq = utr3_dict.get(header, "")
        
        # Remove 5' UTR
        if cdna_seq.startswith(utr5_seq):
            cdna_seq = cdna_seq[len(utr5_seq):]
        
        # Remove 3' UTR
        if cdna_seq.endswith(utr3_seq):
            cdna_seq = cdna_seq[:-len(utr3_seq)]
        
        trimmed_sequences[header] = cdna_seq
    return trimmed_sequences

def write_fasta(output_file, sequences):
    """Write a dictionary of sequences to a FASTA file."""
    with open(output_file, "w") as f:
        for header, seq in sequences.items():
            f.write(f">{header}\n{seq}\n")

def main():
    # Input FASTA file paths
    cdna_file = "cdna.fasta"
    utr5_file = "utr5.fasta"
    utr3_file = "utr3.fasta"
    
    # Output file path
    output_file = "trimmed_cdna.fasta"
    
    # Load FASTA files as dictionaries
    print("Loading input FASTA files...")
    cdna_dict = load_fasta_as_dict(cdna_file)
    utr5_dict = load_fasta_as_dict(utr5_file)
    utr3_dict = load_fasta_as_dict(utr3_file)
    
    # Trim UTRs from cDNA sequences
    print("Trimming UTRs...")
    trimmed_sequences = remove_utrs(cdna_dict, utr5_dict, utr3_dict)
    
    # Write the trimmed sequences to a new FASTA file
    print(f"Writing trimmed sequences to {output_file}...")
    write_fasta(output_file, trimmed_sequences)
    print("Done!")

if __name__ == "__main__":
    main()
```
