## Supplementary Files 1-18 for "The co-receptors *Orco* and *Ir8a* are required for coordinated expression of chemosensory genes in the antennae of the yellow fever mosquito, *Aedes aegypti*": Supplementary 10□ R_scripts.txt

##TxImport.R
#Instructions
#	1. Download results from Salmon job. If run on Galaxy, you will need to change the file type to quant.sf. Place into the working directory.
#	2. Load the tx2gene.csv mapping file. This file links all transcript IDs to their associated Gene IDs for analysis at the gene level. Place into the working directory.
#	3. Set working directory and run the script. This will provide gene level count data.

# Load Required Libraries
if (!requireNamespace("BiocManager", quietly = TRUE)) install.packages("BiocManager")
if (!requireNamespace("tximport", quietly = TRUE)) BiocManager::install("tximport")
if (!requireNamespace("DESeq2", quietly = TRUE)) BiocManager::install("DESeq2")
if (!requireNamespace("readr", quietly = TRUE)) install.packages("readr")
if (!requireNamespace("dplyr", quietly = TRUE)) install.packages("dplyr")

library(tximport)
library(DESeq2)
library(readr)
library(dplyr)

# Set working directory to folder containing quant.sf files
dir <- "~/path/"  # <-- Change this to your actual path

# List all quant.sf files in the directory
files <- list.files(dir, pattern = "quant.sf", full.names = TRUE, recursive = TRUE)


# Assign sample names based on folder names (modify if needed)
sample_names <- basename(files)
names(files) <- sample_names

print(names(files))

# Load Transcript-to-Gene Mapping File
tx2gene <- read.csv("tx2gene.csv", header = TRUE, stringsAsFactors = FALSE)  # MODIFY PATH IF NEEDED


# Import Transcript Quantifications and Summarize to Gene Level
txi <- tximport(files, type = "salmon", tx2gene = tx2gene, countsFromAbundance = "no")

write.csv(txi$counts, "tximport_counts.csv")

##deseq2_analysis.R
#Instructions
#	1. Place results from TxImport into working directory.
#	2. Upload colData.csv into your working directory. This is the metadata that defines which sample belongs to which strain.
#	3. Run the script. you will get the following output: DESeq2_results_orco_vs_orlando.csv; DESeq2_results_ir8a_vs_orlando.csv; volcano_orco_vs_orlando.png; volcano_ir8a_vs_orlando.png
#	4. Note: The DESeq2 results files include comparisons of the each mutant to orlando. The DESeq2 analysis was performed with all three strains, and the results files highlight differences between each mutant and the wildtype.
#	5. Note: The volcano plots are just for an initial visualization to ensure that the script worked as expected.

# Check and install missing packages only if necessary
required_packages <- c("DESeq2", "ggplot2", "RColorBrewer")

new_packages <- required_packages[!(required_packages %in% installed.packages()[,"Package"])]

if(length(new_packages)) install.packages(new_packages)

# Load the necessary libraries
library(DESeq2)
library(ggplot2)
library(RColorBrewer)

# Load the count data and coldata
counts <- read.csv("tximport_counts.csv", row.names = 1)
coldata <- read.csv("coldata.csv", row.names = 1)

# Ensure the count data is rounded to integers (if not already done)
counts <- round(counts)

# Set the row names of colData to match the column names of counts
rownames(coldata) <- colnames(counts)

# Check the data
head(counts)
head(coldata)

# Create the DESeqDataSet object
dds <- DESeqDataSetFromMatrix(countData = counts, colData = coldata, design = ~strain)

# Run DESeq2 analysis
dds <- DESeq(dds)

# Results for Orco vs Orlando
res_orco <- results(dds, contrast = c("strain", "Orco", "Orlando"))
summary(res_orco)

# Results for Ir8a vs Orlando
res_ir8a <- results(dds, contrast = c("strain", "Ir8a", "Orlando"))
summary(res_ir8a)

# Plot volcano plot for Orco vs Orlando
volcano_orco <- ggplot(as.data.frame(res_orco), aes(x = log2FoldChange, y = -log10(pvalue))) +
  geom_point(aes(color = padj < 0.05), size = 2) +
  scale_color_manual(values = c("gray", "red")) +
  theme_minimal() +
  labs(title = "Volcano Plot: Orco vs Orlando", x = "Log2 Fold Change", y = "-Log10(p-value)")

# Plot volcano plot for Ir8a vs Orlando
volcano_ir8a <- ggplot(as.data.frame(res_ir8a), aes(x = log2FoldChange, y = -log10(pvalue))) +
  geom_point(aes(color = padj < 0.05), size = 2) +
  scale_color_manual(values = c("gray", "red")) +
  theme_minimal() +
  labs(title = "Volcano Plot: Ir8a vs Orlando", x = "Log2 Fold Change", y = "-Log10(p-value)")

# Save volcano plots
ggsave("volcano_orco_vs_orlando.png", plot = volcano_orco)
ggsave("volcano_ir8a_vs_orlando.png", plot = volcano_ir8a)

# Export DESeq2 results to CSV
write.csv(as.data.frame(res_orco), "DESeq2_results_orco_vs_orlando.csv")
write.csv(as.data.frame(res_ir8a), "DESeq2_results_ir8a_vs_orlando.csv")

##Filter_DEGs.R
#Instructions
#	1. Load the DESeq2 results files into your working directory. These are the .csv files resulting from deseq2_analysis.R
#	2. You will need to run this script a total of 4 times, twice per DESeq2 output file.
#	3. First load the orco_vs_orlando result file. Set log2FoldChange > 1. Set the name of the output file to DEGs_Orco_UP_salmon.txt. This will return a list of genes that are overexpressed in Orco.
#	4. Next, set log2FoldChange <-1. Set the name of the output file to DEGs_Orco_DN_salmon.txt. This will return a list of genes that are underexpressed in Orco.
#	5. Repeat steps 3 and 4 with Ir8a to obtain a list of genes that are over- or under-expressed in Ir8a.

# Load necessary library
library(dplyr)

# Load DESeq2 results file
deseq_results <- read.table("DESeq2_results_ir8a_vs_orlando.csv", header = TRUE, sep = ",")  # Replace with your file path

# Filter genes based on p-value and log2 fold change thresholds
filtered_genes <- deseq_results %>%
  filter(padj < 0.05 & (log2FoldChange > 1))

# Select only the first column (gene list)
gene_list <- filtered_genes[, 1]

# Display the filtered gene list
print(gene_list)

# Optionally, save the gene list to a new file
write.table(gene_list, "DEGs_Ir8a_UP_salmon.txt", sep = "\t", row.names = FALSE, col.names = FALSE, quote = FALSE)

##volcano_plot_highlight_chemosensory_labelpoints.R
#Instructions
#	1. Place the DESeq2 results files (.csv) into the working directory.
#	2. Place the lists of chemosensory genes (Ors.txt, IRs.txt, GRs.txt) into the working directory.
#	3. Run the script once for Orco and once for Ir8a. Adjust the name of the plot title on line 108 for the respective strains.
#	4. This will generate volcano plots highlighting chemosensory genes, and will label the most significant chemosensory genes at the sides of the plot, ordering the labels from top to bottom in decreasing significance.

library(ggplot2)
library(dplyr)
library(ggrepel)
library(stringr)  # For text manipulation

# Load DESeq2 output
deseq_results <- read.table("DESeq2_results_ir8a_vs_orlando.csv", header = TRUE, sep = ",")  # Replace with your file path

# Load gene lists
OR_genes <- readLines("ORs.txt")  # Replace with your file path
IR_genes <- readLines("IRs.txt")  # Replace with your file path
GR_genes <- readLines("GRs.txt")  # Replace with your file path

# Combine all chemosensory gene lists into a single list
chemosensory_genes <- c(OR_genes, IR_genes, GR_genes)

# Function to determine category based on gene ID
get_category <- function(gene_id) {
  if (is.na(gene_id)) return("Other")  # Handle missing values
  if (gene_id %in% OR_genes || gene_id == "Orco") return("Ors")  # Ensure Orco is red
  if (gene_id %in% IR_genes || gene_id == "Ir8a") return("Irs")  # Ensure Ir8a is orange
  if (gene_id %in% GR_genes) return("Grs")
  return("Other")
}

# Apply the function to create a Category column
deseq_results <- deseq_results %>%
  mutate(Category = sapply(GeneID, get_category)) %>%
  mutate(Significance = ifelse(padj < 0.05 & abs(log2FoldChange) > 1, "Significant", "Not Significant"))

# Assign colors based on category and significance
deseq_results <- deseq_results %>%
  mutate(Color = case_when(
    GeneID == "Orco" ~ "Ors",
    GeneID == "Ir8a" ~ "Irs",
    Significance == "Significant" ~ Category,
    Significance == "Not Significant" ~ "grey80",  # Lighter grey for non-significant genes
    TRUE ~ "Other"
  ))

# Exclude extreme values from being labeled
filtered_deseq_results <- deseq_results %>%
  filter(log2FoldChange > -20 & log2FoldChange < 20)

# Ensure all significantly upregulated chemoreceptors are labeled
upregulated_chemoreceptors <- filtered_deseq_results %>%
  filter(Category %in% c("Ors", "Irs", "Grs") & log2FoldChange > 1 & padj < 0.05)

# Ensure the top 10 significantly downregulated chemoreceptors are labeled
downregulated_chemoreceptors <- filtered_deseq_results %>%
  filter(Category %in% c("Ors", "Irs", "Grs") & log2FoldChange < -1 & padj < 0.05) %>%
  arrange(padj) %>%
  head(10)

# Ensure Orco and Ir8a are always labeled
always_label <- filtered_deseq_results %>%
  filter(GeneID %in% c("Orco", "Ir8a"))

# Combine all labels and compute fixed label x positions
labels <- bind_rows(upregulated_chemoreceptors, downregulated_chemoreceptors, always_label) %>%
  distinct(GeneID, .keep_all = TRUE) %>%
  mutate(ShortLabel = str_remove(GeneID, "_transcript$"),
         label_x = ifelse(log2FoldChange < 0, -10, 10))  # Fixed label position

# **Adjust Y positions to avoid overlap (evenly distribute between 0 and 90)**
labels <- labels %>% arrange(desc(-log10(padj)))  # Sort by significance

# Separate labels into left (-10) and right (+10) groups
left_labels <- labels %>% filter(label_x == -10)
right_labels <- labels %>% filter(label_x == 10)

# Define fixed Y-axis range (0 to 90)
y_min <- 10
y_max <- 90

# Evenly distribute Y positions within the range
left_labels$adjusted_y <- seq(y_max, y_min, length.out = nrow(left_labels))
right_labels$adjusted_y <- seq(y_max, y_min, length.out = nrow(right_labels))

# Recombine labels after adjusting Y positions
labels <- bind_rows(left_labels, right_labels)

# **Plot**
plot <- ggplot() +
  # Scatter plot for all points, excluding extreme values
  geom_point(data = filtered_deseq_results, 
             aes(x = log2FoldChange, y = -log10(padj), color = Color), 
             size = 0.8, alpha = 0.75) +
  # Segments from data points to adjusted label positions
  geom_segment(data = labels, 
               aes(x = log2FoldChange, xend = label_x, 
                   y = -log10(padj), yend = adjusted_y), 
               color = "#808080", size = 0.3) +
  # Labels at fixed -10 or +10 positions with adjusted Y
  geom_label(data = labels, 
             aes(x = label_x, y = adjusted_y, label = ShortLabel),
             size = 3, fill = "white", color = "black", label.size = 0.2, alpha = 0.75) +
  # Axis limits
  scale_x_continuous(limits = c(-15, 15), oob = scales::squish) +
  scale_y_continuous(limits = c(0, 110), oob = scales::squish) +
  # Colors
  scale_color_manual(
    values = c("Ors" = "#FF0000", "Irs" = "#FFAF00", "Grs" = "#FF00FF", 
               "Other" = "grey20", "Not Significant" = "grey90"),  # Grey color for non-significant points
    guide = guide_legend(override.aes = list(alpha = 1, size = 2))
  ) +
  # Titles and theme
  labs(title = expression(italic("Ir8a"^"-/-") ~ "vs Orlando"), 
       x = "log2 Fold Change", 
       y = expression(-log[10](italic(p)[adj]))) +
  theme_minimal() +
  theme(
    legend.title = element_blank(), 
    plot.margin = margin(10, 10, 10, 10),
    panel.background = element_blank(),
    plot.background = element_rect(fill = "white", color = "white"),
    axis.line = element_line(color = "black")
  ) +
  # Threshold lines
  geom_hline(yintercept = -log10(0.05), linetype = "dashed", color = "grey20") +
  geom_vline(xintercept = c(-1, 1), linetype = "dashed", color = "grey20")

# Save the plot
ggsave("Labeled_Volcano_Plot_Updated.png", plot = plot, width = 5.5, height = 5.5, dpi = 600)

# Print the plot
print(plot)

##volcano_plot_TFs.R
#Instructions
#	1. Place the DESeq2 results files (.csv) into the working directory.
#	2. Place the lists of TF genes (TFs_Other.txt, TFs_miR279.txt, TFs_Chemosensory.txt, TFs_MMBdREAM.txt) into the working directory.
#	3. Run the script once for Orco and once for Ir8a. Adjust the name of the plot title on line 108 for the respective strains.
#	4. This will generate volcano plots highlighting TFs, and will label any TFs that don't fall within the TFs_Other.txt category.

# Load required libraries
library(ggplot2)
library(dplyr)
library(ggrepel)  # For improved text positioning

# Load DESeq2 results
deseq_results <- read.table("DESeq2_results_ir8a_vs_orlando.csv", header = TRUE, sep = ",")

# Load TF gene lists
Other_TF_genes <- readLines("TFs_Other.txt")             
miR_279_pathway_genes <- readLines("TFs_miR279.txt")     
Chemosensory_TF_genes <- readLines("TFs_Chemosensory.txt")
MMBdREAM_TF_genes <- readLines("TFs_MMBdREAM.txt")

# Categorize genes
get_category <- function(gene_id) {
  if (is.na(gene_id)) return(NA)
  if (gene_id %in% Other_TF_genes) return("Other_TF")
  if (gene_id %in% miR_279_pathway_genes) return("miR_279")
  if (gene_id %in% Chemosensory_TF_genes) return("Chemosensory_TF")
  if (gene_id %in% MMBdREAM_TF_genes) return("MMBdREAM_TF")
  return("Other")
}

# Apply function
deseq_results <- deseq_results %>%
  mutate(Category = sapply(GeneID, get_category)) %>%
  mutate(Significant = ifelse(padj < 0.05 & abs(log2FoldChange) > 1, "Significant", "Not Significant"))

# Remove rows with NA in Category or Significant
deseq_results <- deseq_results %>%
  filter(!is.na(Category), !is.na(Significant))

# Assign colors for categories
color_palette <- c("MMBdREAM_TF" = "#56B4E9",  # MMB/dREAM complex
                   "miR_279" = "#009E73",    # miR-279 pathway
                   "Chemosensory_TF" = "red",  # Chemosensory TFs
                   "Other_TF" = "#E69F00",  # Other TFs
                   "Not Significant" = "#B0B0B0")  # Grey for Not Significant

# Filter out "Other" category
filtered_deseq_results <- deseq_results %>%
  filter(Category != "Other")

# Select all significant genes
significant_genes <- filtered_deseq_results %>%
  filter(Significant == "Significant")

# Exclude "Other_TF" from labeling
significant_genes <- significant_genes %>%
  filter(Category != "Other_TF")

# Define color logic based on significance
filtered_deseq_results$color_group <- ifelse(filtered_deseq_results$Significant == "Significant", 
                                             filtered_deseq_results$Category, 
                                             "Not Significant")

# Plot
plot <- ggplot(filtered_deseq_results, aes(x = log2FoldChange, y = -log10(padj), color = color_group)) +
  geom_point(size = 0.8, alpha = 0.75) +
  geom_text_repel(data = significant_genes, aes(label = GeneID),
                  color = "black",   # Ensures labels are black
                  box.padding = 0.3, 
                  point.padding = 0.2, 
                  segment.color = "grey50",  # Color for the line segments
                  segment.size = 0.5,  # Adjust the segment line thickness
                  max.overlaps = Inf, 
                  size = 3) +
  scale_color_manual(values = color_palette, 
                     breaks = c("MMBdREAM_TF", "miR_279", "Chemosensory_TF", "Other_TF", "Not Significant"),
                     labels = c("MMB/dREAM complex", "miR-279 pathway", "Chemosensory TFs", "Other TFs", "Not Significant")) +
  scale_x_continuous(limits = c(-5, 5), oob = scales::squish) +
  scale_y_continuous(limits = c(0, 30), oob = scales::squish) +
  labs(title = expression(italic("Ir8a"^"-/-") ~ "vs Orlando Transcription Factors"), 
       x = "log2 Fold Change", 
       y = expression(-log[10](italic(p)[adj]))) +
  theme_minimal() +
  theme(legend.title = element_blank(),
        panel.background = element_blank(),
        plot.background = element_rect(fill = "white", color = "white"),
        axis.line = element_line(color = "black")) +
  geom_hline(yintercept = -log10(0.05), linetype = "dashed", color = "grey20") +
  geom_vline(xintercept = c(-1, 1), linetype = "dashed", color = "grey20") +
  guides(color = guide_legend(override.aes = list(alpha = 1), 
                              keyheight = unit(0.5, "cm"), 
                              keywidth = unit(1.5, "cm")))  # Adjust legend appearance

# Save and print
ggsave("Labeled_Volcano_Plot_Segments.png", plot = plot, width = 5.5, height = 2.5, dpi = 600)
print(plot)


##Heatmaps_ComplexHeatmap.R
#Instructions
#	1. From the Salmon quant.sf output files, locate the column with the header TPM.
#	2. Copy that header into a new excel file.
#	3. Do this for all samples. Average the TPM values for all samples within each strain (exclude Orlando 4).
#	4. Make sure the headers are as follows: Gene_Name	Orlando	Orco	IR8a.
#	5. Save as a tabular file named TPMs_Salmon.txt. Place in working directory.
#	6. Place ORs.txt, IRs.txt, and GRs.txt into the working directory.
#	7. Run the script. This will produce a heatmap that displays each chemoreceptor family as a separate heatmap and combines them all into one figure. This only shows the top 80 genes, where at least one strain meets a threshold for plotting. The number of genes plotted can be adjusted altering the numbers on lines 38 and 39.

# Install and load the necessary packages
if (!requireNamespace("ComplexHeatmap", quietly = TRUE)) install.packages("ComplexHeatmap")
if (!requireNamespace("gridExtra", quietly = TRUE)) install.packages("gridExtra")
if (!requireNamespace("dplyr", quietly = TRUE)) install.packages("dplyr")
library(dplyr)
library(ComplexHeatmap)
library(gridExtra)

# Sample TPM data (replace with your actual data)
tpm_data <- read.table("TPM_data_file_Salmon.txt", header = TRUE, sep = "\t")  # Replace with your file path

# Gene lists for ORs, IRs, and GRs
OR_genes <- readLines("ORs.txt")  # Replace with your file path
IR_genes <- readLines("IRs.txt")  # Replace with your file path
GR_genes <- readLines("GRs.txt")  # Replace with your file path

# Combine all gene lists into a single list
gene_lists <- list(OR = OR_genes, IR = IR_genes, GR = GR_genes)

# Define color palettes for each gene family
color_schemes <- list(
  OR = colorRampPalette(c("black", "#FF9999", "#FF0000"))(100),
  IR = colorRampPalette(c("black", "#FFCC99", "#FFAF00"))(100),
  GR = colorRampPalette(c("black", "#FFCCFF", "#FF00FF"))(100)
)

# Apply log10 transformation to TPM data (log10(TPM+1))
tpm_data_transformed <- tpm_data %>%
  mutate(across(-Gene_Name, ~log10(. + 1)))  # Transform TPM values, keeping the Gene_Name column unchanged

# Reorder the columns to ensure Orlando is between Orco and IR8a
column_order <- c("Orco", "Orlando", "IR8a", setdiff(colnames(tpm_data_transformed), c("Orco", "Orlando", "IR8a", "Gene_Name")))
tpm_data_transformed <- tpm_data_transformed[, c("Gene_Name", column_order)]

# Function to determine cutoff for each gene type to retain 50 genes
determine_cutoff <- function(filtered_data) {
  sorted_values <- sort(apply(filtered_data[, c("Orco", "Orlando", "IR8a")], 1, max), decreasing = TRUE)
  if (length(sorted_values) >= 80) {
    return(sorted_values[80])  # 50th highest value as cutoff
  } else {
    return(min(sorted_values))  # If fewer than 50 genes, use the minimum max-expression value
  }
}

# Function to plot heatmap with ComplexHeatmap
plot_complex_heatmap <- function(gene_list, data, title, color_scheme) {
  # Filter the data for genes in the current list
  filtered_data <- data %>% filter(Gene_Name %in% gene_list)
  
  # Sort rows by TPM values in the Orlando column
  filtered_data <- filtered_data %>% arrange(desc(Orlando))
  
  # Apply cutoff based on max expression in key columns
  cutoff <- determine_cutoff(filtered_data)
  filtered_data <- filtered_data %>% filter(apply(.[, c("Orco", "Orlando", "IR8a")], 1, max) >= cutoff)
  
  # Set up the matrix for plotting (rows = genes, columns = samples)
  data_matrix <- as.matrix(filtered_data[, -1])  # Exclude Gene_Name column for the heatmap matrix
  
  # Set Gene_Name as row names
  rownames(data_matrix) <- filtered_data$Gene_Name
  
  # Create a heatmap using ComplexHeatmap
  ht <- Heatmap(data_matrix,
                name = title,  # Title for the heatmap
                col = color_scheme,  # Apply the specific color scheme
                cluster_rows = FALSE,  # No clustering of rows
                cluster_columns = FALSE,  # No clustering of columns
                show_row_names = TRUE,  # Show row (gene) names
                show_column_names = TRUE,  # Show column (sample) names
                row_names_gp = gpar(fontsize = 8),  # Adjust font size for row names (genes)
                column_names_gp = gpar(fontsize = 9),  # Adjust font size for column names (samples)
                column_names_side = "top",  # Place column labels at the top
                column_names_centered = TRUE,
                column_names_rot = 0,  # Ensure column labels are not tilted
                column_title = title,  # Place labels at the top
                heatmap_legend_param = list(
                  title = expression(log[10] (TPM + 1)),  # Subscript 10
                  direction = "horizontal"
                ))
  
  return(ht)
}

# Create heatmaps for each gene family
ht_or <- plot_complex_heatmap(gene_lists$OR, tpm_data_transformed, "Odorant Receptors", color_schemes$OR)
ht_ir <- plot_complex_heatmap(gene_lists$IR, tpm_data_transformed, "Ionotropic Receptors", color_schemes$IR)
ht_gr <- plot_complex_heatmap(gene_lists$GR, tpm_data_transformed, "Gustatory Receptors", color_schemes$GR)

# Concatenate heatmaps horizontally
ht_list = ht_or + ht_ir + ht_gr
# Dynamically adjust height while ensuring it does not exceed 9 inches
plot_height <- min(9.5, 0.6 * nrow(tpm_data_transformed) * 0.5)  # Calculate height based on number of genes, capped at 9 inches

# Save the heatmap as a PNG file with width 5.5 inches and dynamic height
png("Heatmaps.png", width = 5.5, height = 8.3, units = "in", res = 600)
draw(ht_list, auto_adjust = FALSE, heatmap_legend_side = "bottom", 
     padding = unit(c(2, 5, 2, 2), "mm"))  # Increase left margin to 6mm
dev.off()

## Final_topGO.R
#Instructions
#	1. Place the Final_AGWG_GOterms_no_transcript.gaf file into the working directory.
#	2. Place the four .csv files that are lists of DEGs (from Filter_DEGs.R) into the working directory.
#	3. Run the script. This will result in GO term enrichment results .csv files, as well as a plot showing the most enriched GO terms as a .png file.
#	4. Run the script for each list of DEGs. Change the name of the title for the plot on line 118 to accurately describe what is being plotted for each list of DEGs.

# Install and Load Required Packages
if (!requireNamespace("BiocManager", quietly = TRUE)) install.packages("BiocManager")
if (!requireNamespace("topGO", quietly = TRUE)) BiocManager::install("topGO")
if (!requireNamespace("GO.db", quietly = TRUE)) BiocManager::install("GO.db")

library(topGO)
library(GO.db)
library(ggplot2)
library(scales)
library(stringr)  # Load stringr for text wrapping

# Read the GAF file (assuming it’s in the current directory)
gaf_file <- "Final_AGWG_GOterms_no_transcript.gaf"
gaf_data <- read.delim(gzfile(gaf_file), header = FALSE, comment.char = "!", sep = "\t")

# Map Genes to GO Terms (assumes columns 2 and 5 contain gene ID and GO term)
geneID2GO <- with(gaf_data, tapply(V5, V2, function(x) as.character(unique(x))))

# Load list of differentially expressed genes (DEGs)
interesting_genes <- readLines("DEGs_Orco_DN_salmon.txt")

# Remove suffixes (e.g., '-RA' or any other two-letter suffix)
interesting_genes <- gsub("-[A-Z]{2}$", "", interesting_genes)

# Define a gene universe
all_genes <- unique(names(geneID2GO))

# Define a named factor indicating which genes are of interest
gene_list <- factor(as.integer(all_genes %in% interesting_genes))
names(gene_list) <- all_genes

# Create a topGOdata object
GOdata <- new("topGOdata",
              ontology = "MF",  # Change to BP or CC if needed
              allGenes = gene_list,
              nodeSize = 10,
              annot = annFUN.gene2GO,
              gene2GO = geneID2GO)

# Run GO enrichment test
resultFisher <- runTest(GOdata, algorithm = "classic", statistic = "fisher")

# Extract all p-values
fisher_scores <- score(resultFisher)

# Generate results table
all_results <- GenTable(GOdata, classicFisher = resultFisher, topNodes = length(usedGO(GOdata)))

# Manually assign correct p-values to avoid <1e-30 issue
all_results$classicFisher <- fisher_scores[all_results$GO.ID]
all_results$classicFisher <- as.numeric(gsub("<", "", all_results$classicFisher))

# Define significance threshold
pval_threshold <- 0.005

# Filter significant results
significant_results <- subset(all_results, classicFisher <= pval_threshold)

# Fixing the "Term" column using GO.db
significant_results$Term <- sapply(significant_results$GO.ID, function(go) {
  Term(go)
})

# Calculate the number of significant genes for each GO term
significant_results$Significant_Genes_Count <- sapply(significant_results$GO.ID, function(go_id) {
  term_genes <- genesInTerm(GOdata, go_id)[[1]]
  significant_genes <- term_genes[term_genes %in% interesting_genes]
  length(significant_genes)
})

# Create the final output dataframe
final_output <- significant_results[, c("GO.ID", "Term", "Annotated", "Significant", "Expected", "classicFisher", "Significant_Genes_Count")]

# Save results to CSV files
write.csv(final_output, "GO_Term_Enrichment_Results.csv", row.names = FALSE)
write.csv(significant_results, "GO_Enrichment_Summary.csv", row.names = FALSE)

# Calculate the log10 of the p-value for ordering
final_output$log10Fisher <- -log10(as.numeric(final_output$classicFisher))

# Reorder the terms such that the smallest p-value (most significant) is at the top
final_output$Term <- factor(final_output$Term, levels = final_output$Term[order(final_output$log10Fisher)])

# Calculate breaks for the color gradient (log10-transformed p-values)
breaks_values <- seq(min(final_output$log10Fisher), max(final_output$log10Fisher), length.out = 6)

# Apply str_wrap() with a reduced width to allow more line breaks
final_output$Term <- str_wrap(final_output$Term, width = 35)  # Increase wrapping for more line breaks

# Generate bar plot with adjusted bar width and reduced text size
plot <- ggplot(final_output, aes(x = reorder(Term, log10Fisher), y = Significant_Genes_Count, fill = log10Fisher)) +
  geom_bar(stat = "identity") +  
  scale_fill_gradient(
    low = "blue", 
    high = "red", 
    name = "p-value",  
    breaks = seq(min(final_output$log10Fisher), max(final_output$log10Fisher), length.out = 6),  
    labels = function(x) format(10^(-x), scientific = TRUE, digits = 2)  # Adjust to show 1 digit after the decimal
  ) +
  coord_flip() +  
  theme_bw() +  
  theme(
    panel.border = element_blank(),  
    legend.position = "bottom",  
    legend.direction = "horizontal",  
    legend.title = element_text(size = 8),  
    legend.text = element_text(size = 8),  # Increase the text size of the legend
    legend.key.height = unit(0.5, "cm"),  # Adjust height for better spacing
    legend.key.width = unit(1.2, "cm"),   # Increase the width of the legend keys
    legend.margin = margin(t = 0, r = 0, b = 0, l = -10),  # Move legend left
    plot.margin = margin(2, 2, 2, 10, "pt"),
    plot.title = element_text(hjust = 2, size = 9),  
    plot.subtitle = element_text(hjust = 0.5, size = 8),  
    axis.text.y = element_text(size = 8),  
    axis.text.x = element_text(size = 8)  
  ) +
  labs(
    title = expression("Functions of genes with decreased abundance in Orco mutant antennae"),
    x = "GO Terms",
    y = "Number of Significant Genes"
  )

# Dynamically adjust width while ensuring the height doesn't exceed 9 inches
plot_width <- min(8, 0.6 * nrow(final_output))  # Adjust width based on the number of GO terms, max width 8 inches

# Save the plot with width of 5.5 inches and height up to 10 inches
ggsave("GO_Term_Enrichment_Plot.png", width = 5.5, height = min(8, 0.4 * nrow(final_output)), units = "in")

##Venn_Diagram.R
#Instructions
#	1. Place Irs.txt, Grs.txt, Ors.txt, All_DEGs.txt, and TFs.txt into the working directory. These are tabular files where each column is a list of all the DEGs (or subsets of DEGs defined by the categories) for the two strains. The headers are as follows: Orco Down	Orco Up	Ir8a Down	Ir8a Up,
#	2. Run the script once for each tabular file. Change the name of the Venn Diagram on line 71 for each category.
#	3. This will result in 4 way Venn Diagram comparisons of genes that are differentially expressed in mutants compared to wildtype.

library(VennDiagram)
library(grid)

# Load the tab-separated file while preserving spaces in column names
deg_data <- read.delim("TFs.txt", header = TRUE, stringsAsFactors = FALSE, na.strings = c("", "NA"), check.names = FALSE)

# Extract column names dynamically (preserve spaces)
list_names <- colnames(deg_data)

# Convert columns to lists while handling missing values
deg_lists <- lapply(deg_data, function(x) na.omit(x[x != ""]))

# Assign lists to variables dynamically
for (i in seq_along(list_names)) {
  assign(list_names[i], deg_lists[[i]], envir = .GlobalEnv)
}

# Define output file
png("venn_diagram.png", width = 3590, height = 3590, res = 600)

# Generate Venn diagram (without category labels)
venn.plot <- draw.quad.venn(
  area1 = length(get(list_names[1])),
  area2 = length(get(list_names[2])),
  area3 = length(get(list_names[3])),
  area4 = length(get(list_names[4])),
  n12 = length(intersect(get(list_names[1]), get(list_names[2]))),
  n13 = length(intersect(get(list_names[1]), get(list_names[3]))),
  n14 = length(intersect(get(list_names[1]), get(list_names[4]))),
  n23 = length(intersect(get(list_names[2]), get(list_names[3]))),
  n24 = length(intersect(get(list_names[2]), get(list_names[4]))),
  n34 = length(intersect(get(list_names[3]), get(list_names[4]))),
  n123 = length(Reduce(intersect, list(get(list_names[1]), get(list_names[2]), get(list_names[3])))),
  n124 = length(Reduce(intersect, list(get(list_names[1]), get(list_names[2]), get(list_names[4])))),
  n134 = length(Reduce(intersect, list(get(list_names[1]), get(list_names[3]), get(list_names[4])))),
  n234 = length(Reduce(intersect, list(get(list_names[2]), get(list_names[3]), get(list_names[4])))),
  n1234 = length(Reduce(intersect, list(get(list_names[1]), get(list_names[2]), get(list_names[3]), get(list_names[4])))),
  category = rep("", 4),  # Hide default category labels
  fill = c("#E69F00", "#56B4E9", "#009E73", "#CC79A7"),
  alpha = 0.5,
  cex = 1.8,
  cat.cex = 1.35,
  fontface = "bold",
  cat.fontface = "bold",
  fontfamily = "sans",
  cat.fontfamily = "sans",
  ind = FALSE,
  margin = 0.08
)

grid.newpage()
grid.draw(venn.plot)

# Define custom labels with italicized first word and superscripted "-/-" without space
formatted_labels <- sapply(list_names, function(name) {
  words <- unlist(strsplit(name, " ", fixed = TRUE))
  if (length(words) >= 2) {
    bquote(italic(.(words[1])) * .(bquote(phantom()^italic("-/-"))) ~ .(words[2]))  
  } else {
    words[1]  # If only one word, keep as is
  }
})

# Manually position labels near the top apex of each oval
grid.text(formatted_labels[[1]], x = 0.12, y = 0.75, gp = gpar(fontsize = 18, fontface = "bold"))
grid.text(formatted_labels[[2]], x = 0.88, y = 0.75, gp = gpar(fontsize = 18, fontface = "bold"))
grid.text(formatted_labels[[3]], x = 0.25, y = 0.85, gp = gpar(fontsize = 18, fontface = "bold"))
grid.text(formatted_labels[[4]], x = 0.75, y = 0.85, gp = gpar(fontsize = 18, fontface = "bold"))

# Add a title
grid.text("Transcription Factors", x = 0.5, y = 0.95, gp = gpar(fontsize = 20, fontface = "bold"))

dev.off()
