## Supplementary Files 1-18 for "The co-receptors *Orco* and *Ir8a* are required for coordinated expression of chemosensory genes in the antennae of the yellow fever mosquito, *Aedes aegypti*": Supplementary 11□ extract_upstream_sequences.py

```
from Bio import SeqIO
import gffutils
import re

# File paths
genome_file = "genome.fasta"  # Path to the genome FASTA file
gff_file = "annotations.gff"  # Path to the GFF file

# Load the genome sequence
genome = SeqIO.to_dict(SeqIO.parse(genome_file, "fasta"))

# Load the GFF file and create a database
db = gffutils.create_db(gff_file, dbfn='annotations.db', force=True, keep_order=True, merge_strategy='merge', sort_attribute_values=True)

# Define the upstream region size
upstream_size = 2000

# Dictionary to store the most upstream TSS per gene
gene_tss = {}

# Loop through each mRNA feature in the GFF file
for feature in db.features_of_type('mRNA'):  # Adjust 'mRNA' to match the feature type in your GFF file
    transcript_id = feature.id
    match = re.match(r'(.+?)(-[A-Za-z0-9]+|_transcript)?$', transcript_id)  # Extract gene ID before the transcript suffix
    gene_id = match.group(1) if match else transcript_id
    chrom = feature.seqid
    strand = feature.strand
    
    # Identify the first CDS element (most upstream CDS) within the mRNA feature
    cds_features = list(db.children(feature, featuretype='CDS'))  # Get all CDS elements within the mRNA feature
    if not cds_features:
        continue  # Skip mRNA if no CDS is found
    
    # Sort the CDS elements by start position (for positive strand) or by end position (for negative strand)
    if strand == '+':
        first_cds = min(cds_features, key=lambda x: x.start)
        tss = first_cds.start  # TSS is the start of the first CDS
    else:
        first_cds = max(cds_features, key=lambda x: x.end)
        tss = first_cds.end  # TSS is the end of the first CDS on the negative strand
    
    # Check if this is the most upstream TSS for the gene
    if gene_id not in gene_tss or (strand == '+' and tss < gene_tss[gene_id]["tss"]) or (strand == '-' and tss > gene_tss[gene_id]["tss"]):
        gene_tss[gene_id] = {"tss": tss, "chrom": chrom, "strand": strand}

# Open a file to save the upstream sequences in FASTA format
with open("upstream_sequences.fasta", "w") as out_file:
    for gene_id, info in gene_tss.items():
        chrom = info["chrom"]
        strand = info["strand"]
        tss = info["tss"]
        
        if strand == '+':
            # For the positive strand, extract upstream sequence
            upstream_start = max(tss - upstream_size, 0)
            upstream_end = tss - 1
            upstream_seq = genome[chrom].seq[upstream_start:upstream_end]
        else:  # For the reverse strand, extract upstream sequence
            upstream_start = tss  # Start just after the TSS
            upstream_end = tss + upstream_size  # Go upstream from the TSS
            upstream_seq = genome[chrom].seq[upstream_start:upstream_end].reverse_complement()
        
        # Write the sequence in FASTA format
        out_file.write(f">{gene_id}\n")
        out_file.write(str(upstream_seq) + "\n")
```
