## Supplementary Files 1-18 for "The co-receptors *Orco* and *Ir8a* are required for coordinated expression of chemosensory genes in the antennae of the yellow fever mosquito, *Aedes aegypti*": Supplementary 12□ Figures S1 - S7.docx

Supplementary Figures

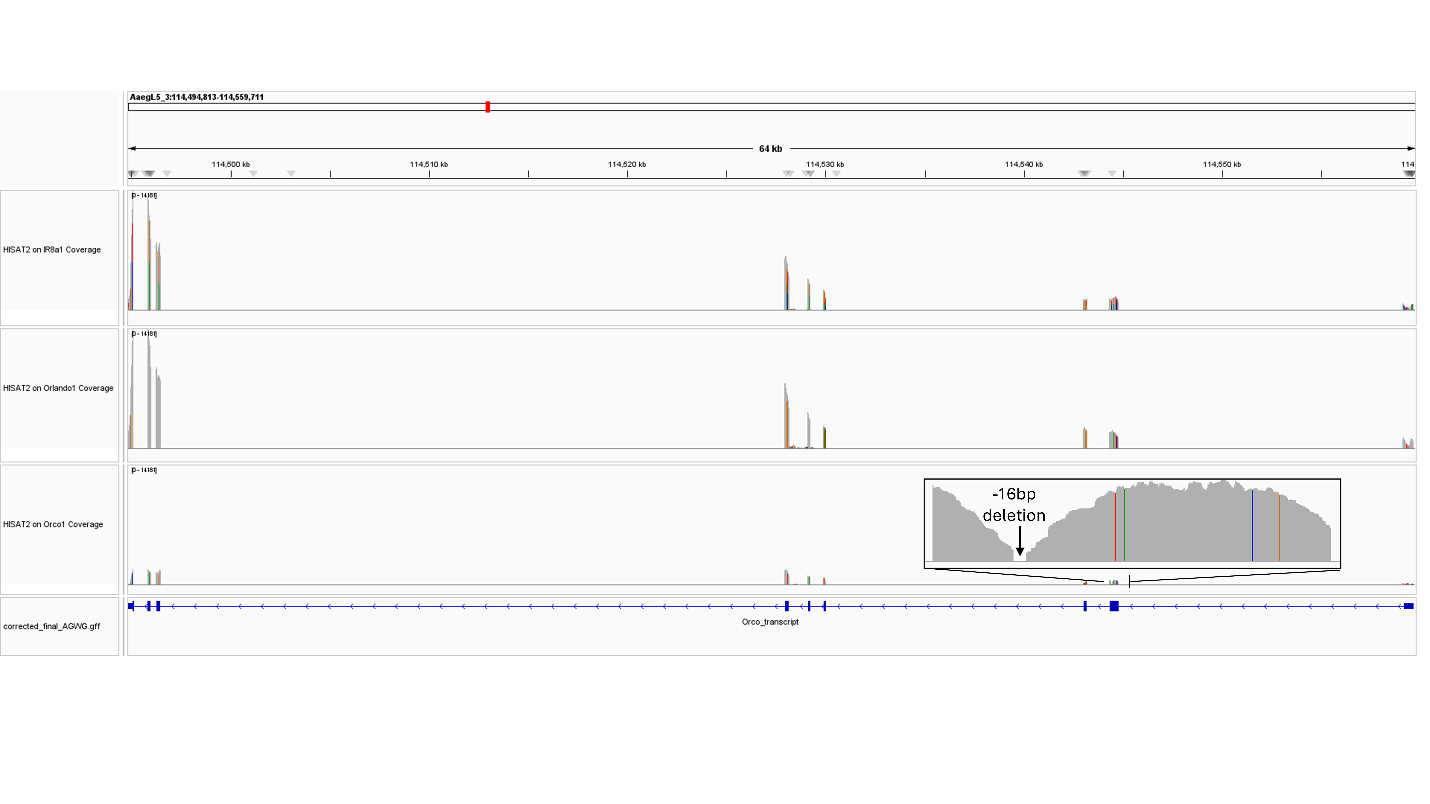

Figure S1. RNAseq coverage of the Orco gene, with the Ir8a^-/-^ strain on top, the Orlando strain in the middle, and the Orco^-/-^ strain on the bottom. Reads aligned with HISAT2 against the *Aedes aegypti* L5 genome. Coverage visualized with Integrated Genome Viewer.

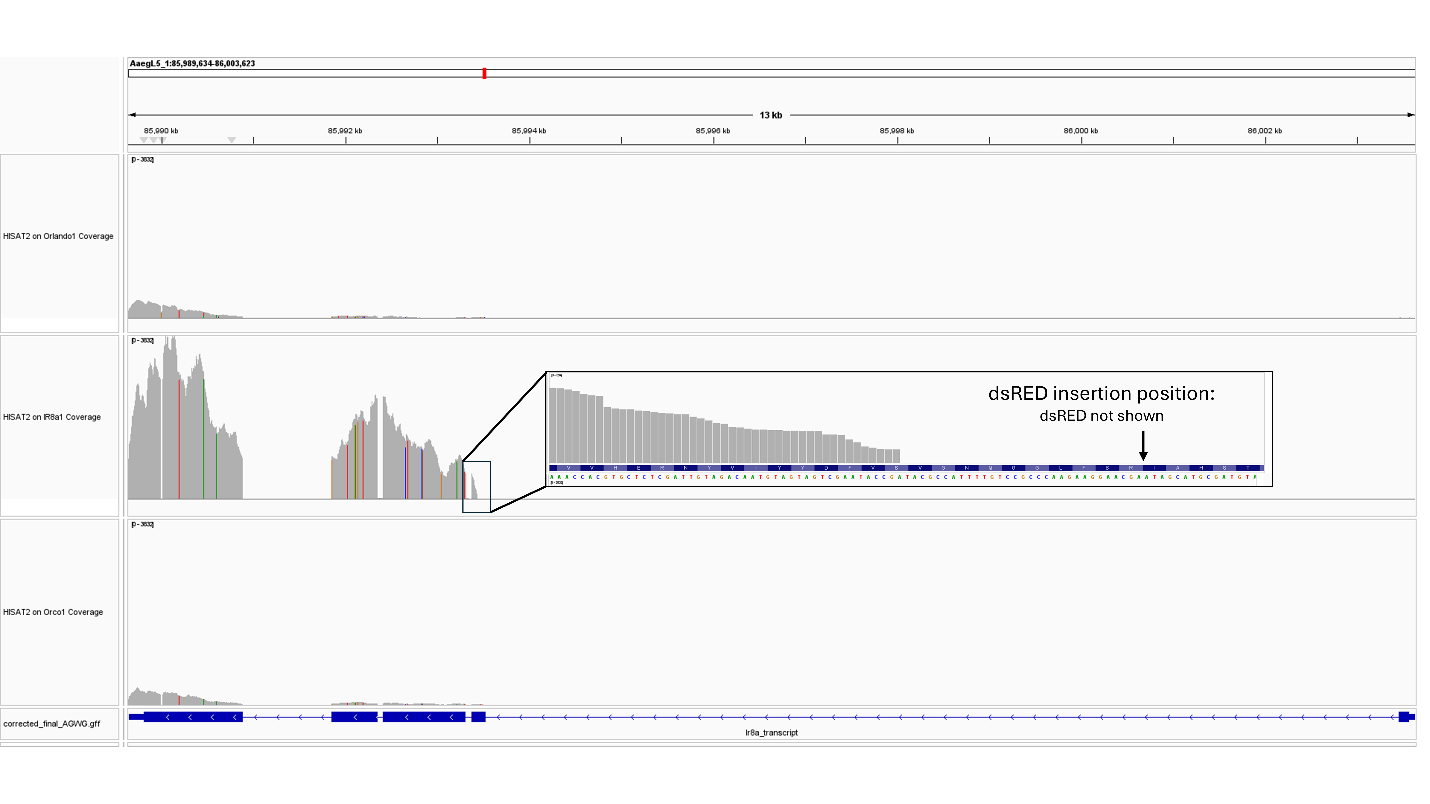

Figure S2. RNAseq coverage of the Ir8a gene, with the Ir8a^-/-^ strain on top, the Orlando strain in the middle, and the Orco^-/-^ strain on the bottom. Reads aligned with HISAT2 against the *Aedes aegypti* L5 genome. Coverage visualized with Integrated Genome Viewer.

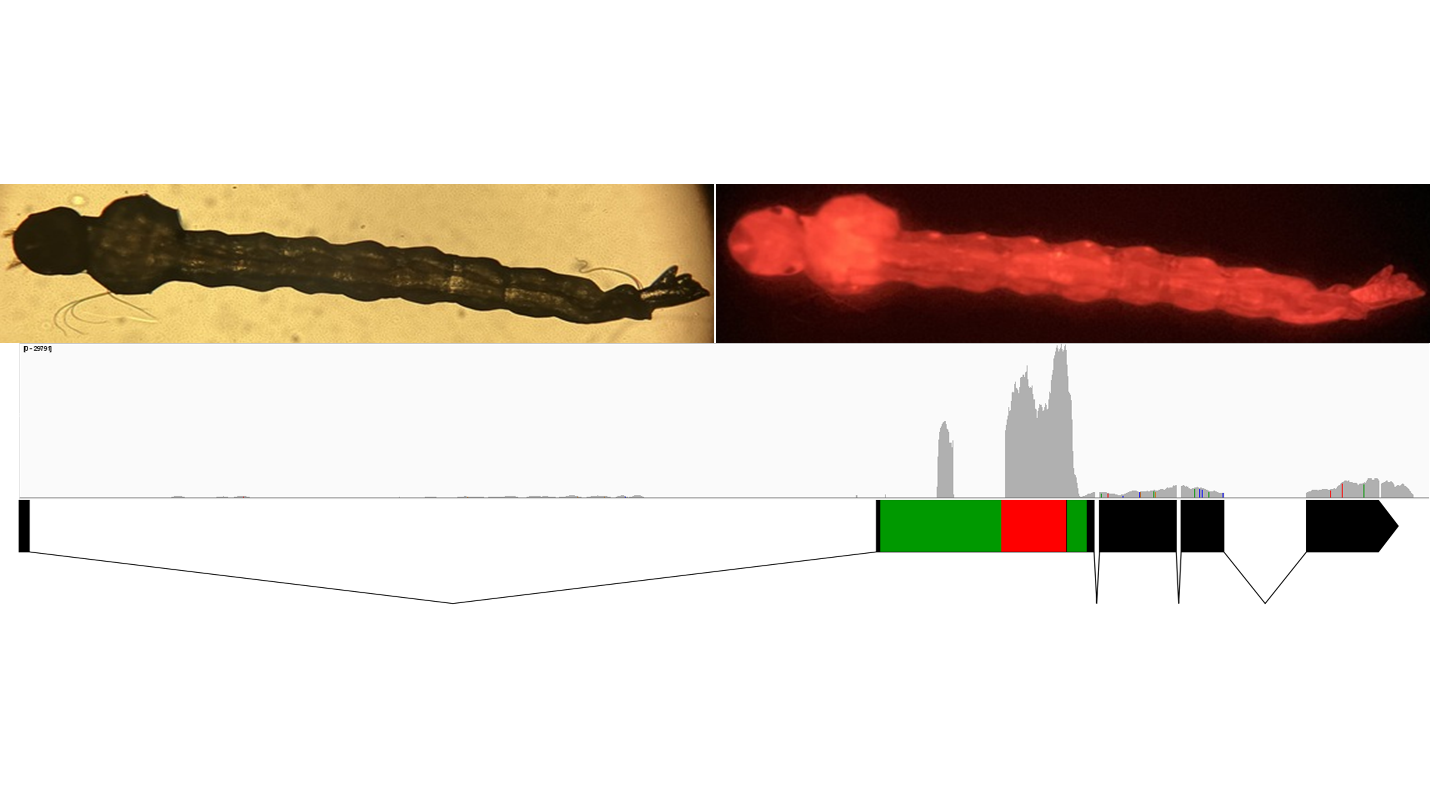
Figure S3. Fluorescent imaging (excitation 558 nm, emission 583 nm) shows that the Ir8a^-/-^ strain of mosquitoes express dsRED in all tissues of the body. HISAT2 alignment was used to align reads from the Ir8a^-/-^ mutant against the Ir8a_dsRED gene, which was set as the “genome”. Coverage was visualized with the Integrated Genome Viewer. A gene map shows the insert in exon 2 (left to right: green = polyubiquitin promoter, red = dsRED ORF, green = SV40). Increased coverage is observed in exons 3-5, following the polyubiquitin promoter.

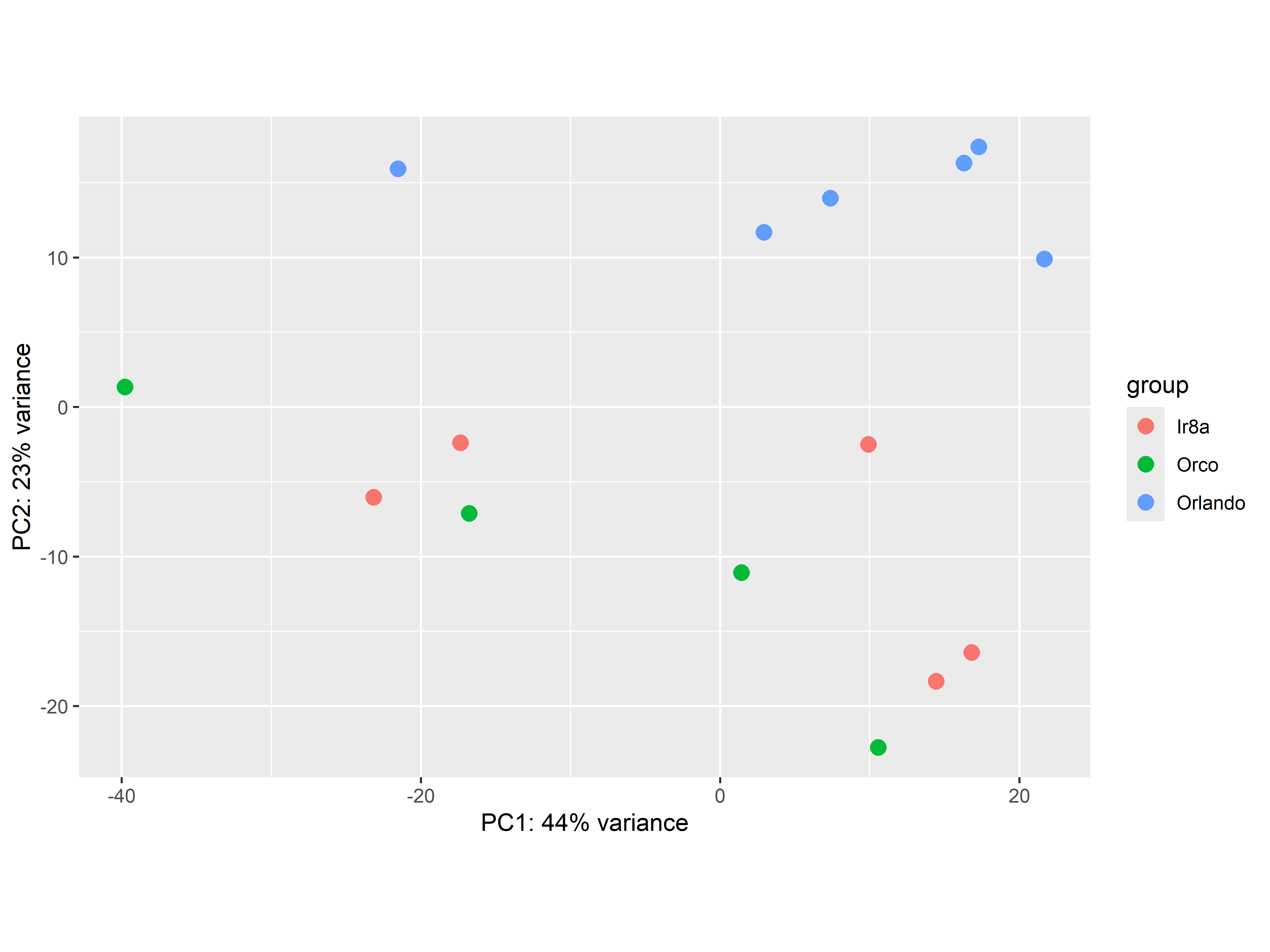

Figure S4. PCA plot of DESeq2 results for all strains shows separation of both mutants from the Orlando wildtype strain, with little separation between mutants.

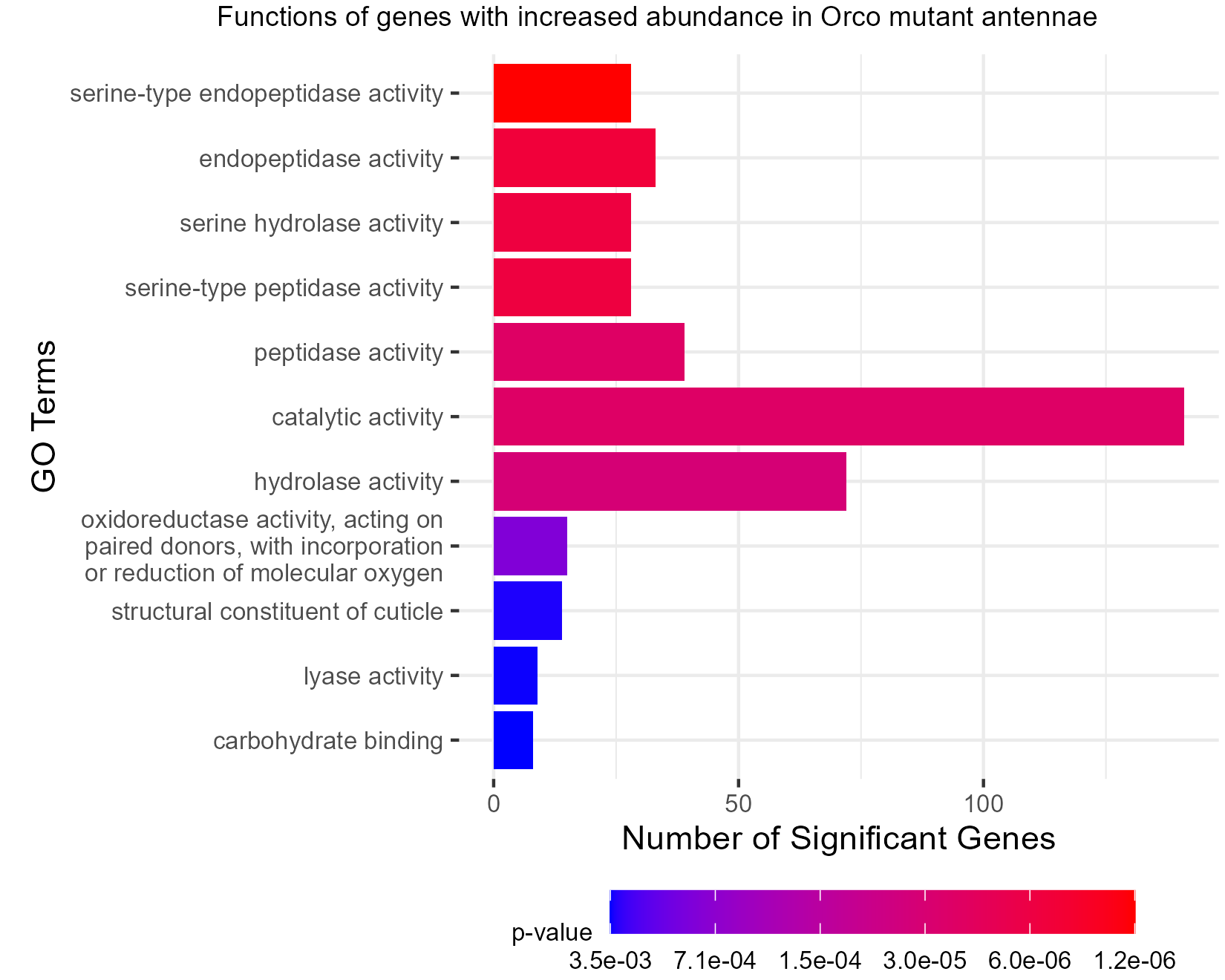

Figure S5. TopGO analysis shows that peptidase functions are representative of the over-expressed genes in the Orco mutant antennae. TopGO analysis was performed at the Molecular Functions level, and the updated GAF file (including the most recently updated chemoreceptor genes) was used to map GO terms to genes. All GO terms with a classic Fisher’s p-value <0.005 were plotted. The color map from red to blue represents the p-value for the enrichment of each term, with the lowest p-values colored in red.

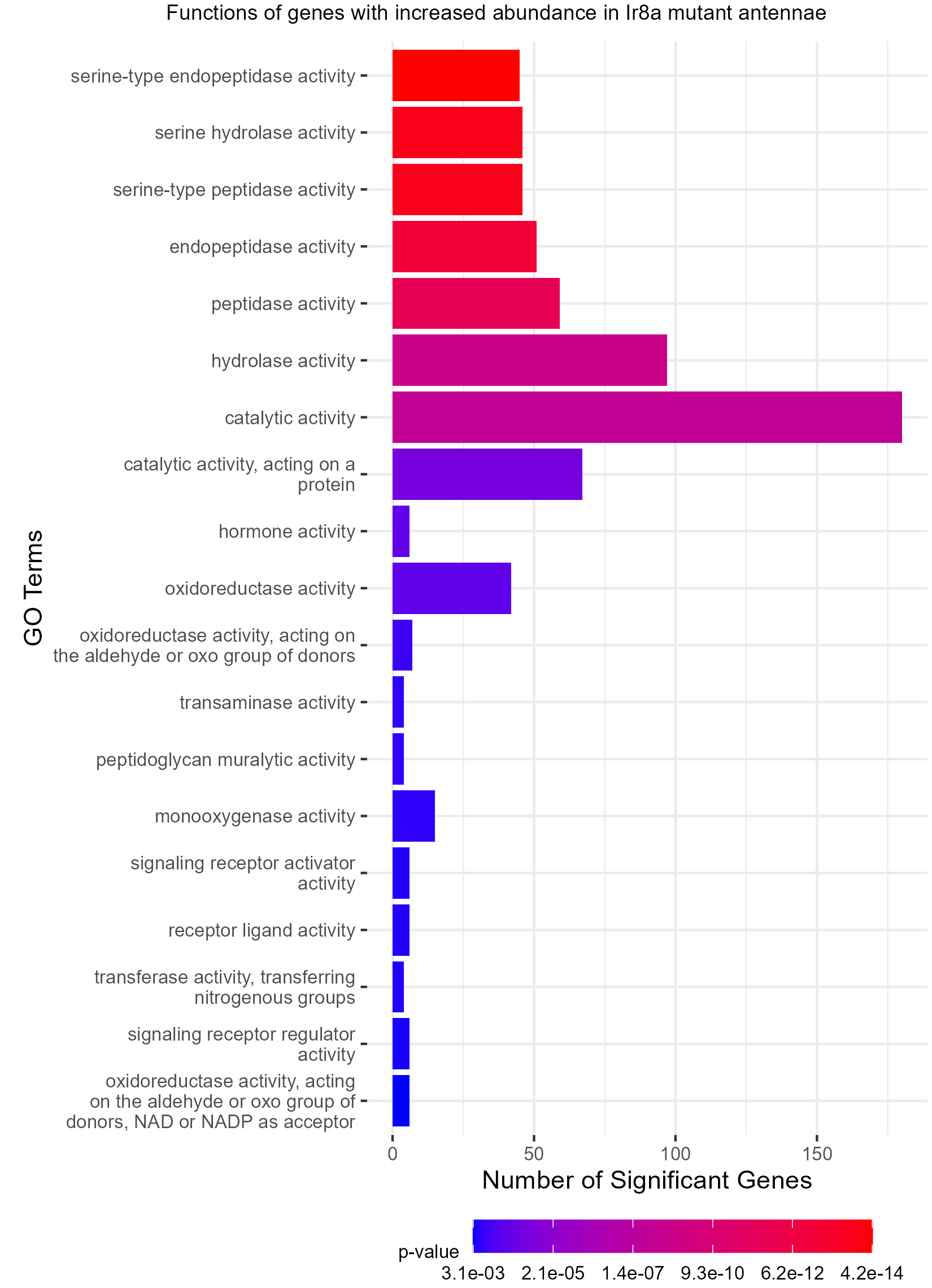

Figure S6. TopGO analysis shows that peptidase functions are representative of the over-expressed genes in the Ir8a mutant antennae. TopGO analysis was performed at the Molecular Functions level, and the updated GAF file (including the most recently updated chemoreceptor genes) was used to map GO terms to genes. All GO terms with a classic Fisher’s p-value <0.005 were plotted. The color map from red to blue represents the p-value for the enrichment of each term, with the lowest p-values colored in red.

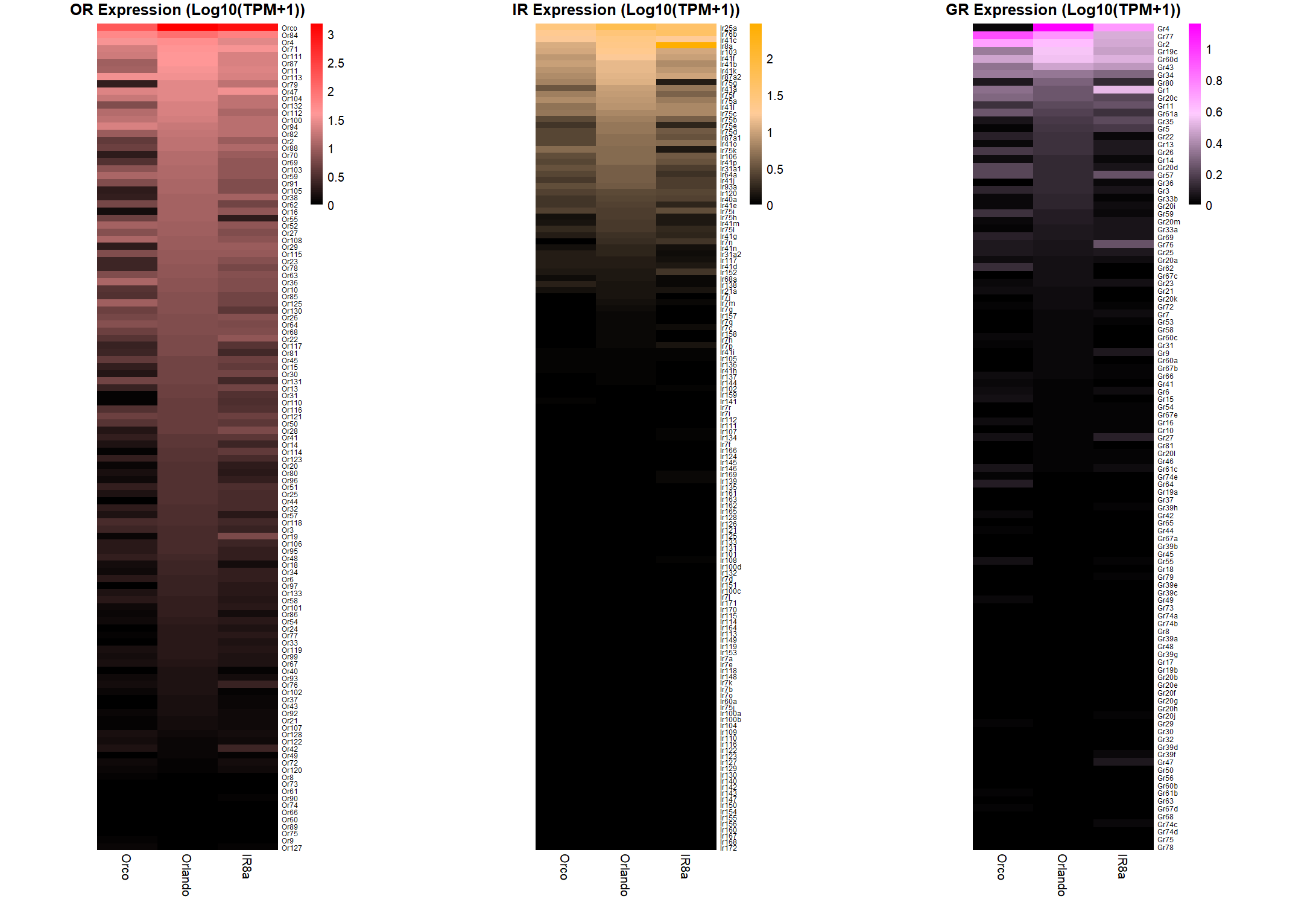

Figure S7. Expanded heatmaps show relative expression levels between strains for different chemoreceptor classes.
