## Supplementary XSTREME Motif Analysis for "The co-receptors *Orco* and *Ir8a* are required for coordinated expression of chemosensory genes in the antennae of the yellow fever mosquito, *Aedes aegypti*": fimo.html

FIMO Results


---

|  |  |  |  |  |  |
| --- | --- | --- | --- | --- | --- |
| **Database and Motifs** | **High-scoring Motif Occurences** | **Debugging Information** | **Results in TSV Format** | **Results in GFF3 Format** | **Best Site per Sequence** |

  
  


---
