## Supplementary XSTREME Motif Analysis for "The co-receptors *Orco* and *Ir8a* are required for coordinated expression of chemosensory genes in the antennae of the yellow fever mosquito, *Aedes aegypti*": sea.html

SEA results


[close ]

[close ]

[
close ]

[
close ]

[
close ]

[
close ]

[
close ]

[
close ]

[
close ]

[
close ]

[
close ]

[
close ]

[
close ]

[
close ]

[
close ]

[
close ]

[
close ]

### SEA

#### Simple Enrichment Analysis

For further information on how to interpret these results please access
https://meme-suite.org/meme/doc/sea-output-format.html.  
To get a copy of the MEME software please access
https://meme-suite.org.

Enriched Motifs
  |  
Input Files
  |  
Program information
  |  
Results in TSV Format 

|  
Matching Sequences

|  
Matching Sites


### Javascript is required to view these results!


#### Enriched Motifs

| Logo | Database | ID | Alt ID | *P*-value | *E*-value | Q-value | TP | FP | Enrichment Ratio | Score Threshold |
| --- | --- | --- | --- | --- | --- | --- | --- | --- | --- | --- |

#### Input Files

###### Alphabet


###### Sequences

###### Motifs

| Database | Source | Motif Count |
| --- | --- | --- |

###### Other Settings

|  |  |
| --- | --- |
| Strand Handling | This alphabet only has one strand. Only the given strand is processed. Both the given and reverse complement strands are processed. |
| Objective Function |  |
| Statistical Test |  |
| Sequence Shuffling |  |
| Hold-out Set |  |
| Pseudocount |  |
| Significance threshold |  |
| Random Number Seed |  |
| Trimming of Control Sequences |  |

###### SEA version

(Release date: )

###### Command line summary
