## Supplementary XSTREME Motif Analysis for "The co-receptors *Orco* and *Ir8a* are required for coordinated expression of chemosensory genes in the antennae of the yellow fever mosquito, *Aedes aegypti*": xstreme.html

XSTREME Results


[close ]

[close ]

[
close ]

[
close ]

[
close ]

[
close ]

[
close ]

[
close ]

[
close ]

[
close ]

[
close ]

[
close ]

### XSTREME

#### Motif Discovery and Enrichment Analysis

For further information on how to interpret these results please access
https://meme-suite.org/meme/doc/xstreme-output-format.html.  
To get a copy of the MEME software please access
https://meme-suite.org.

Motifs
  |  
Programs
  |  
Input Files
  |  
Program information
  |  
Summary in TSV Format 
  |  
Non-redundant Motifs in MEME Text Format


### Javascript is required to view these results!

### Your browser does not support canvas!


#### Motifs

**Enriched motifs
(E-value ≤ ).**

Expand All Clusters
Collapse All Clusters

#### Programs

#### Input Files

###### Alphabet


###### Motifs

###### XSTREME version

(Release date: )

###### Command line
