## Supplementary figures and images for "The co-receptors *Orco* and *Ir8a* are required for coordinated expression of chemosensory genes in the antennae of the yellow fever mosquito, *Aedes aegypti*"

### Figure S1

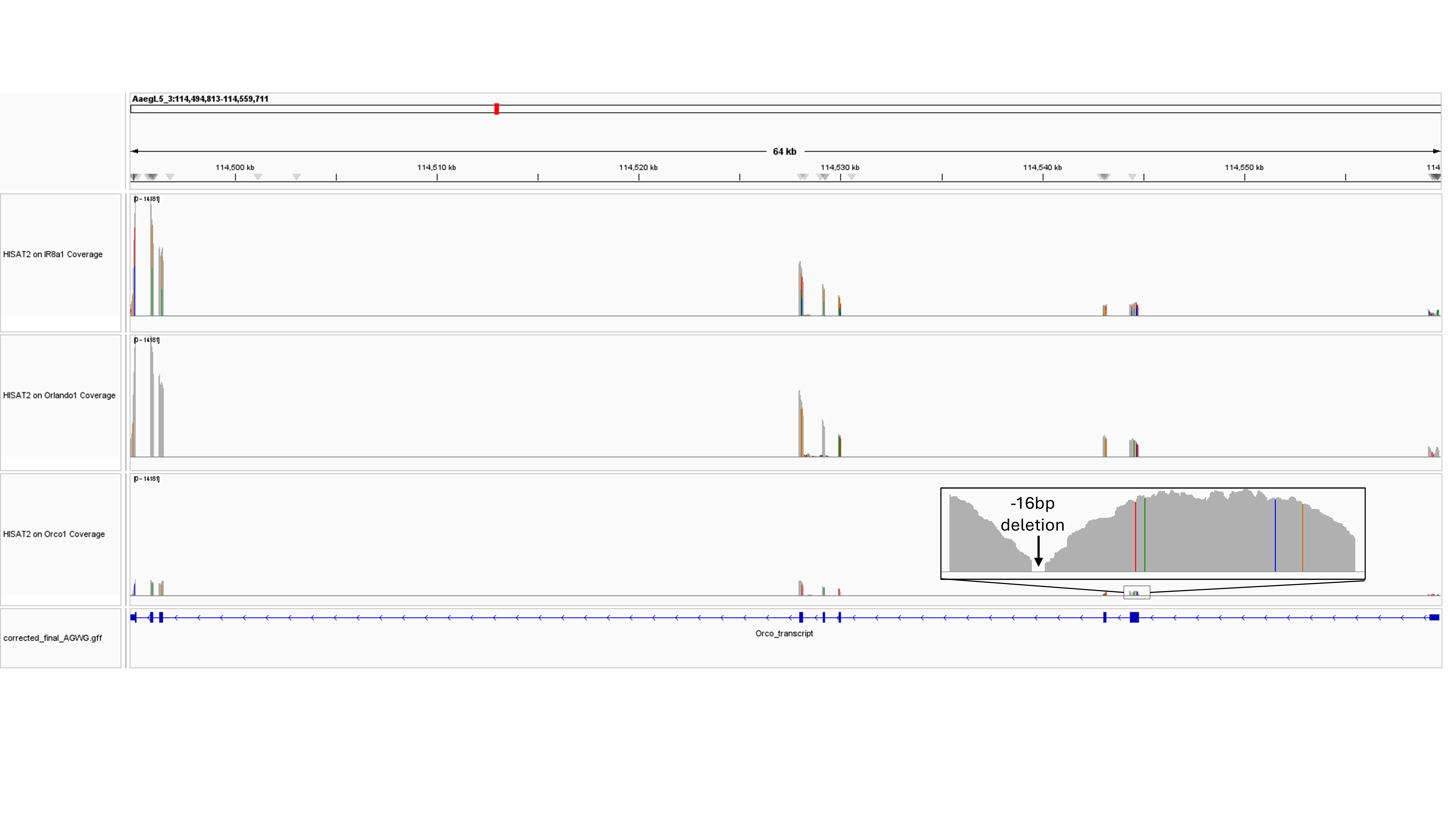

### Figure S2

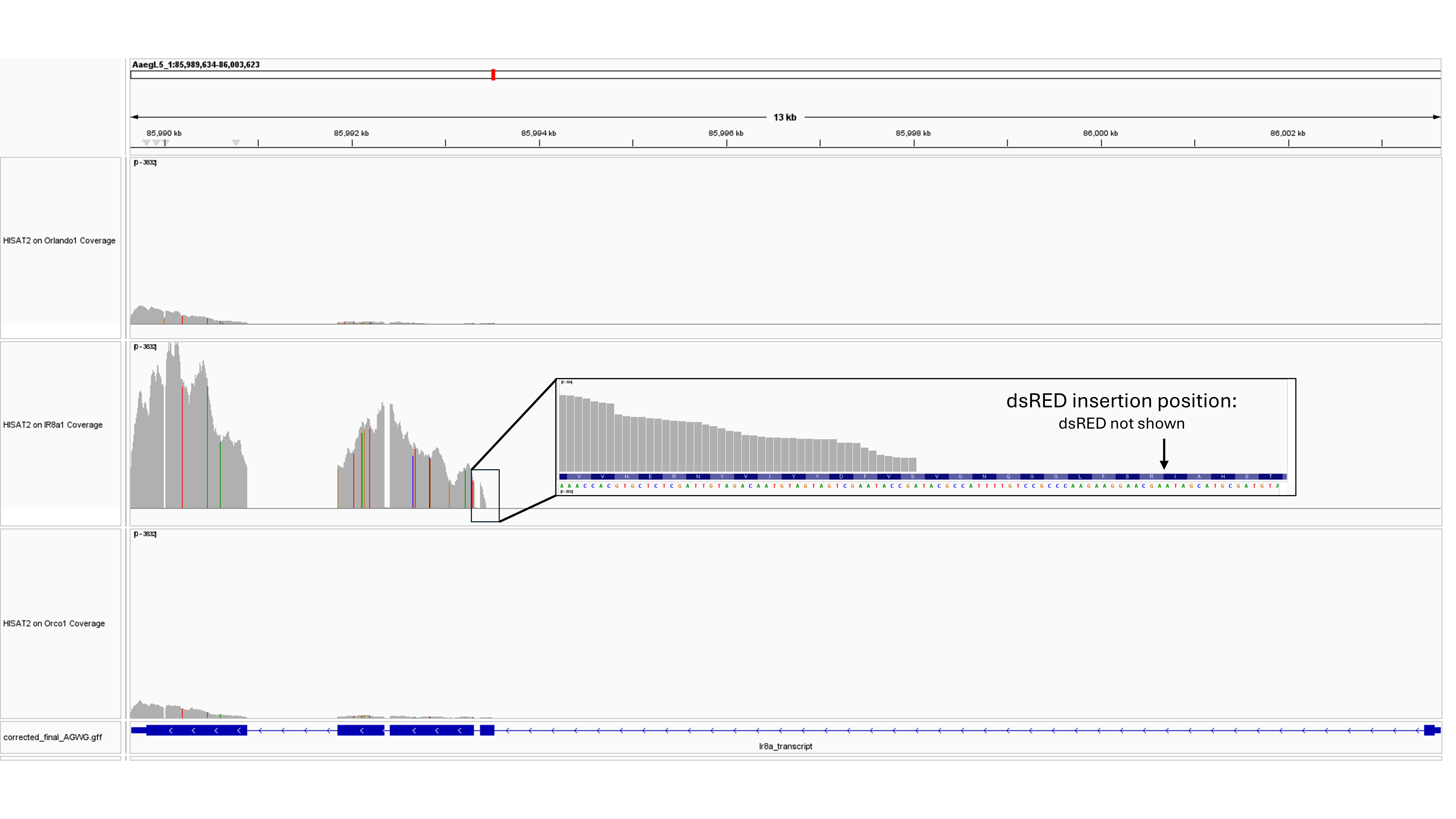

### Figure S3

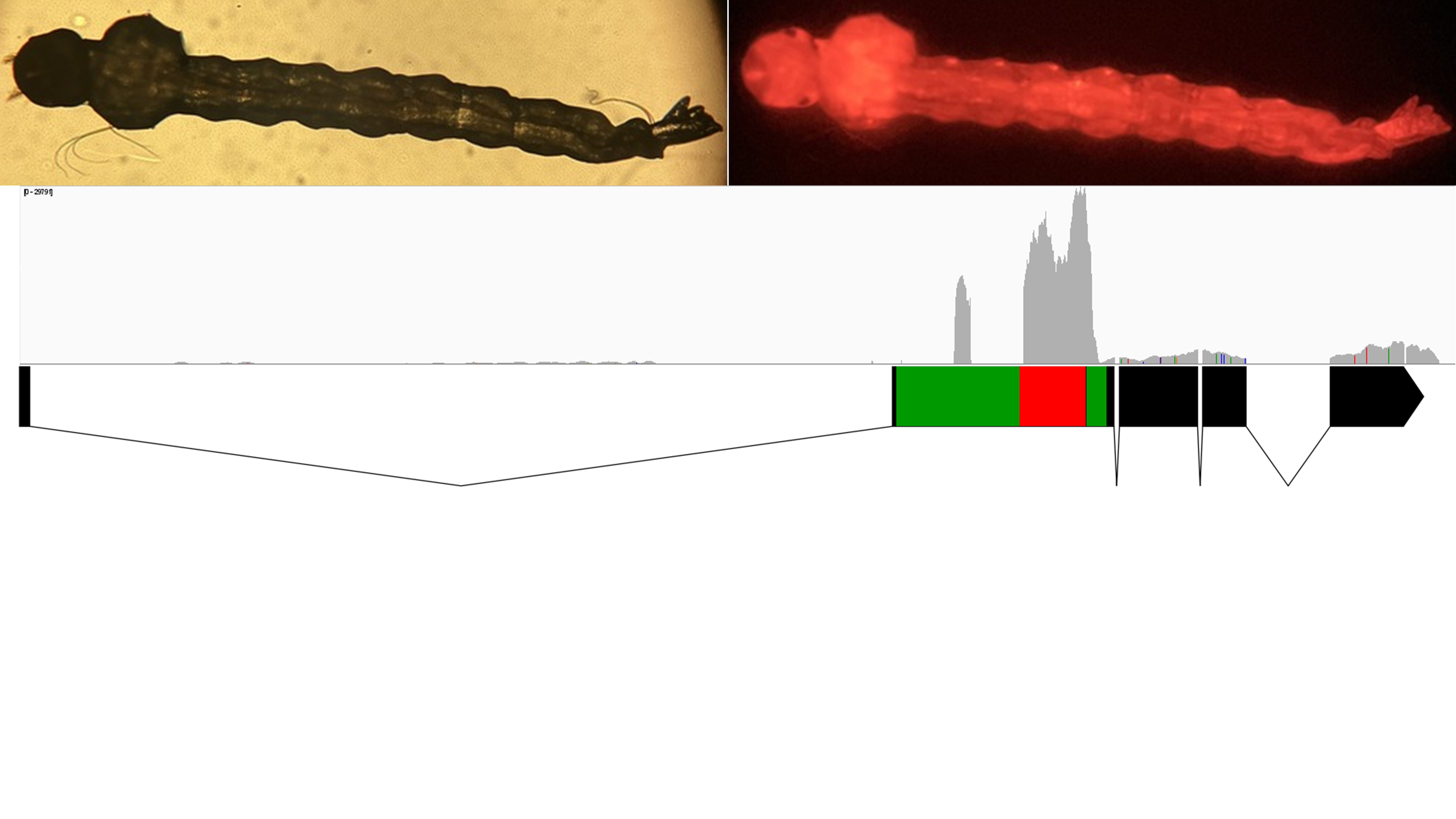

### Figure S4

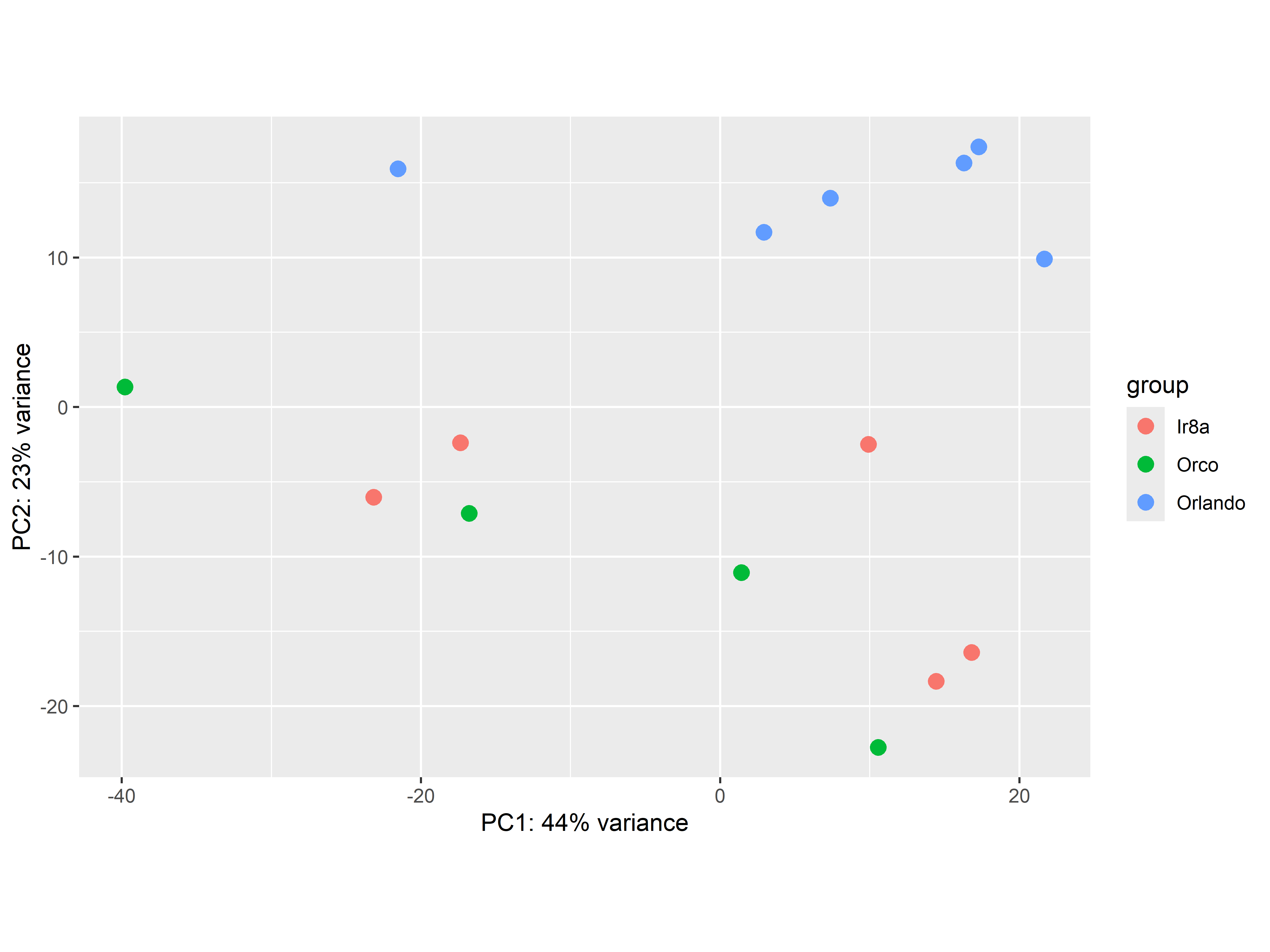

### Figure S5

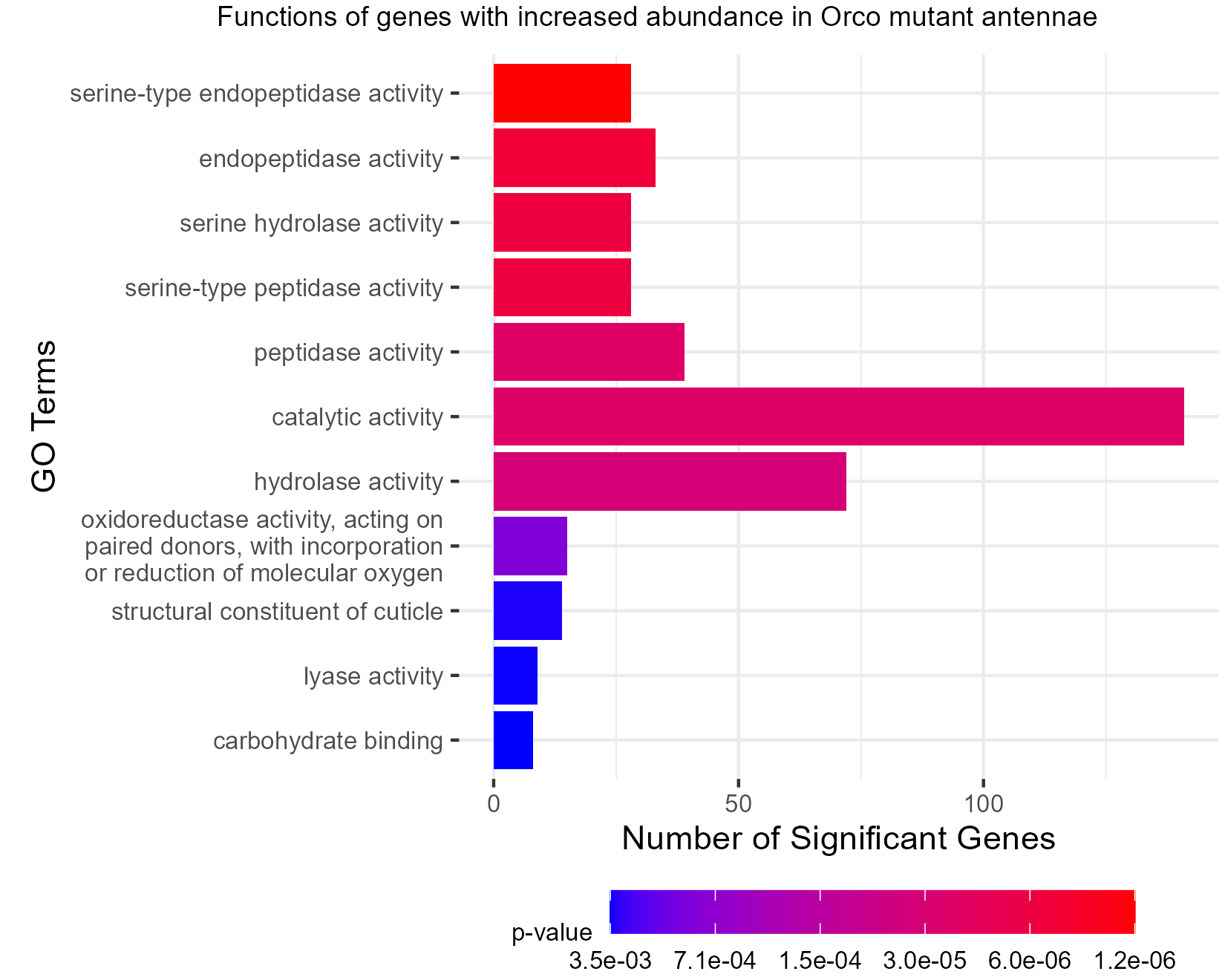

### Figure S6

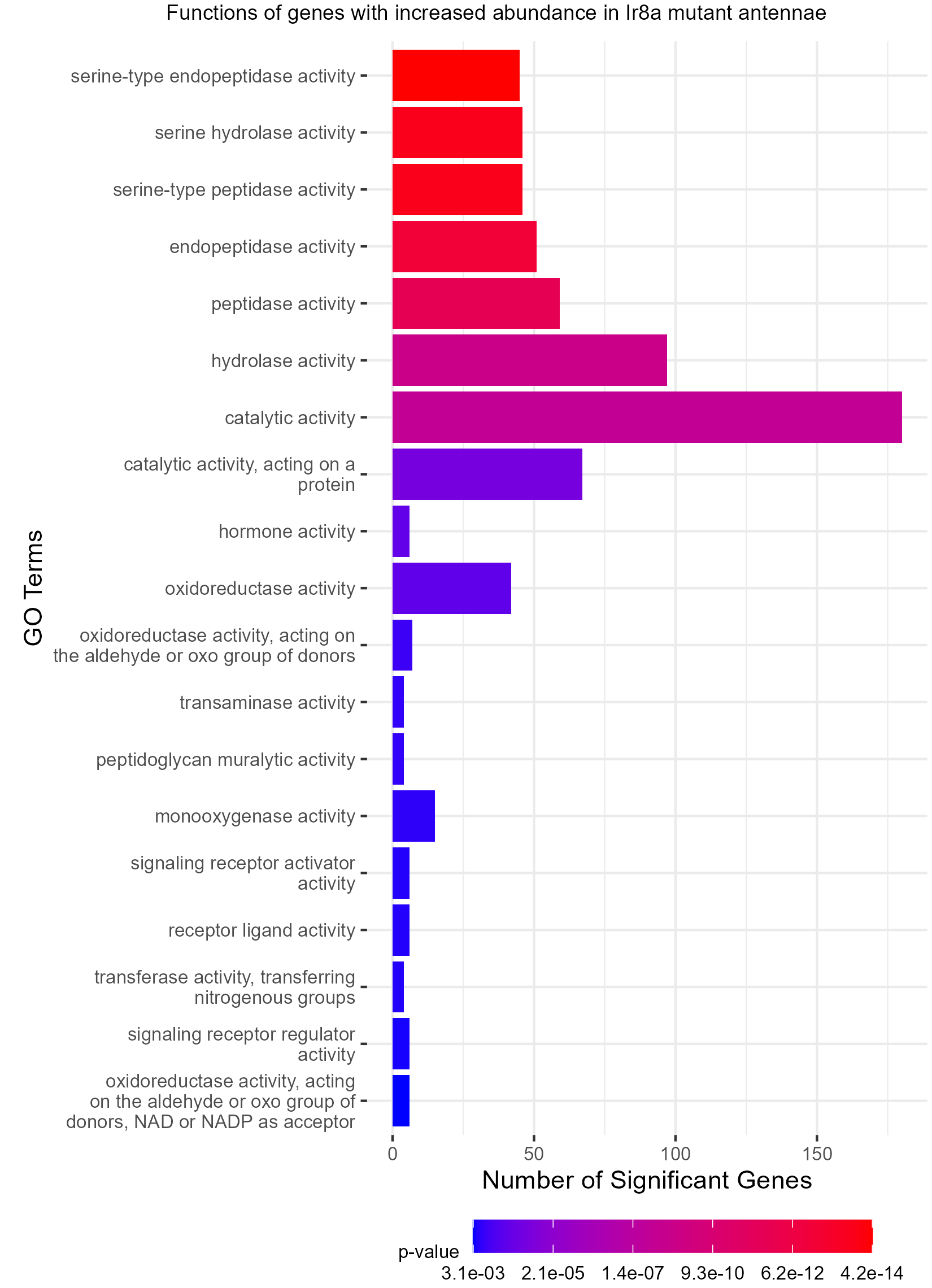

### Figure S7

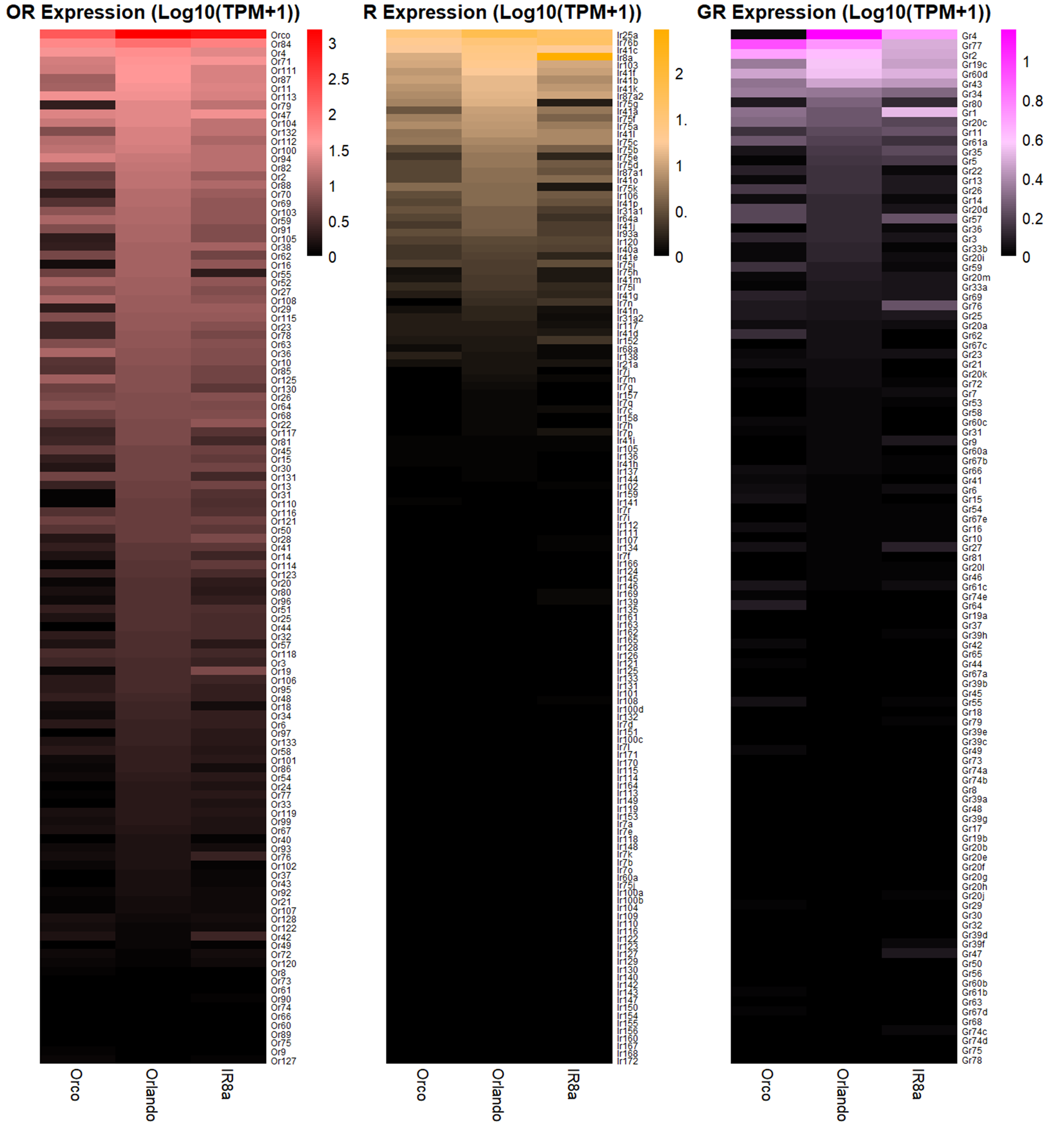

### Figure S8

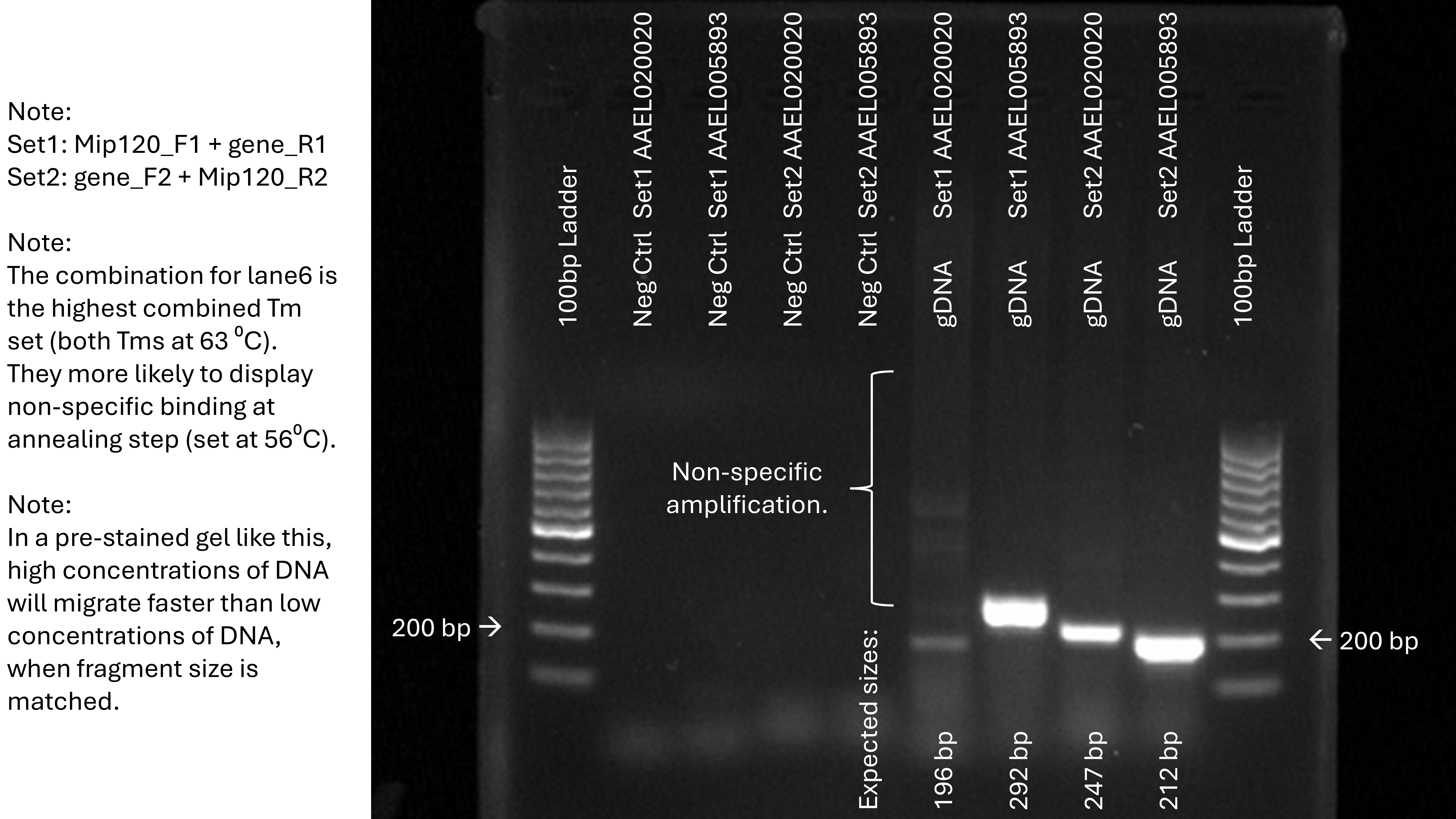

### logo1.png

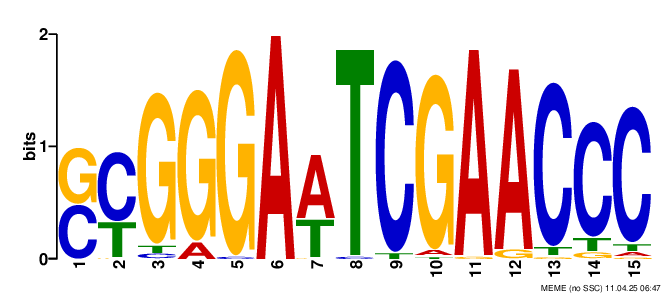

### logo2.png

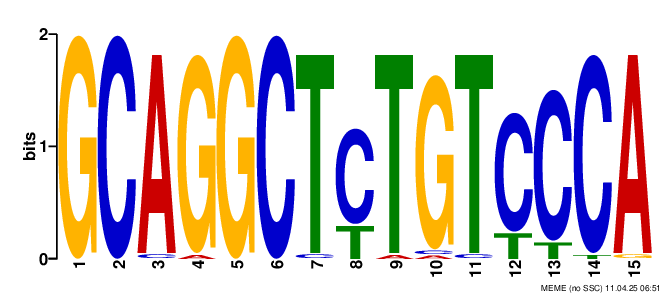

### logo3.png

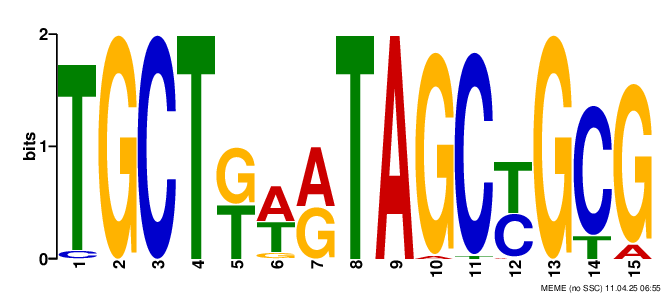

### logo4.png

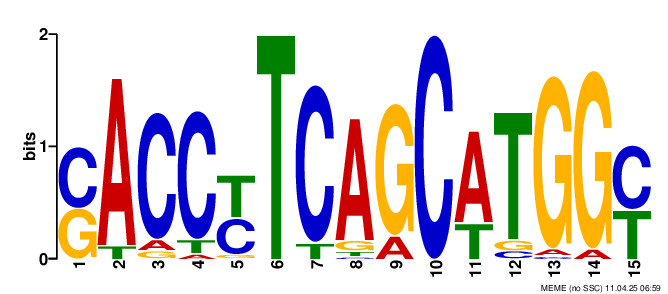

### logo5.png

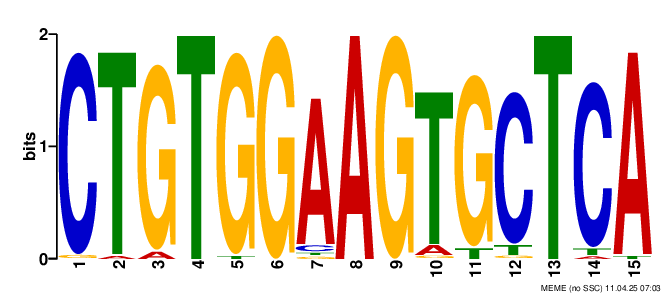

### logo6.png

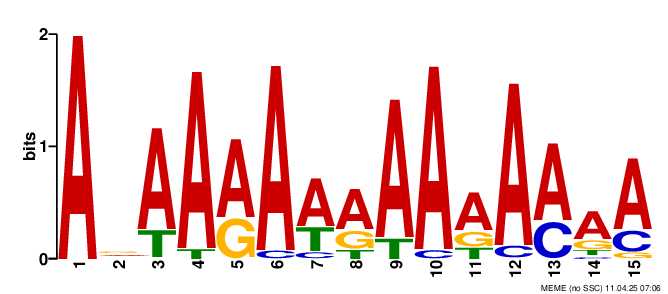

### logo7.png

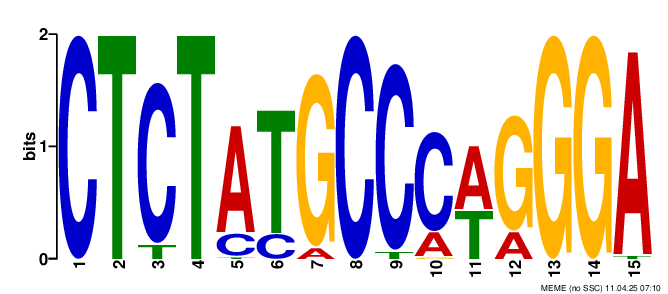

### logo8.png

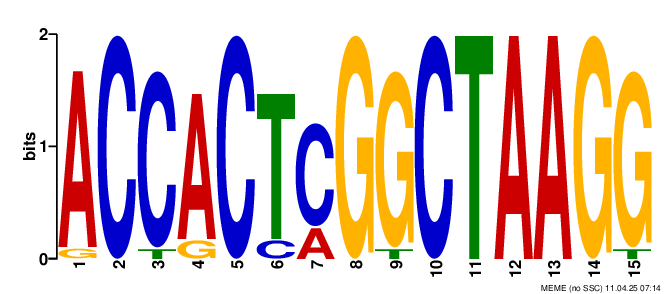

### logo_rc1.png

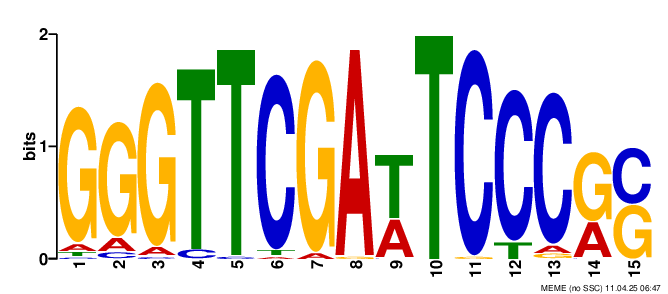

### logo_rc2.png

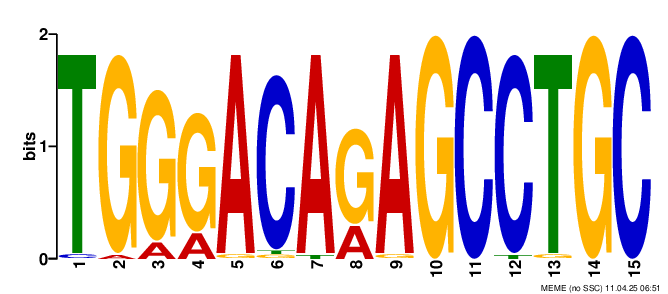

### logo_rc3.png

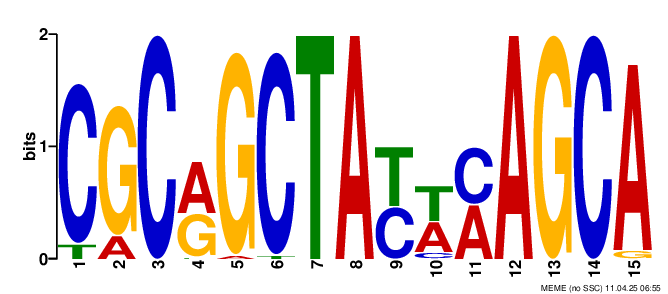

### logo_rc4.png

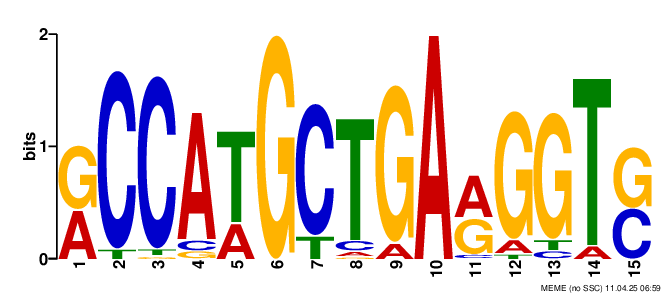

### logo_rc5.png

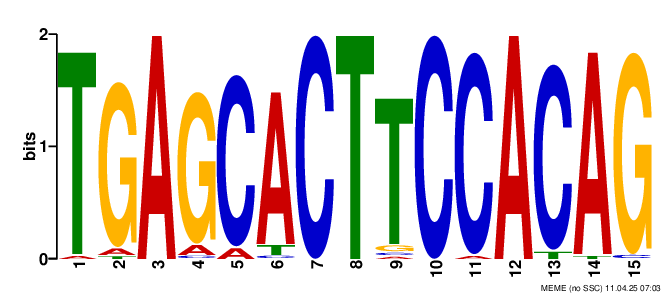

### logo_rc6.png

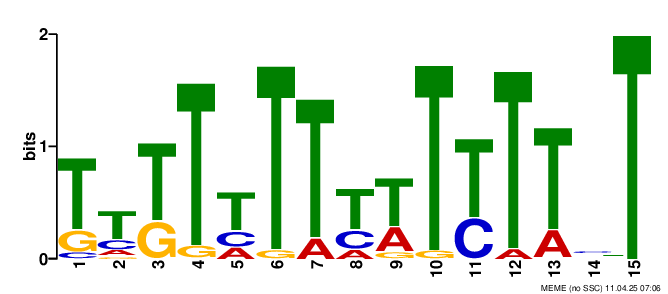

### logo_rc7.png

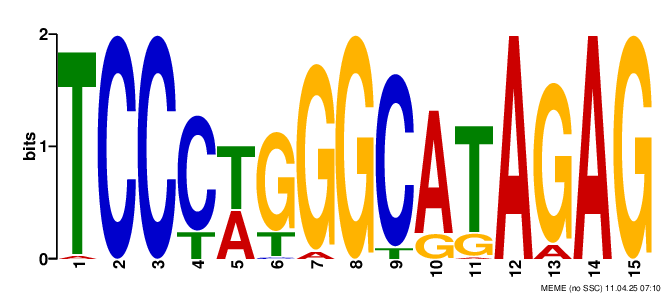

### logo_rc8.png

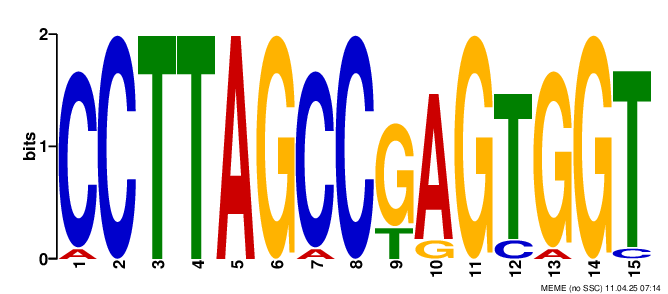
